# A signal peptide-guided platform for in situ functionalization of bacterial nanocellulose in *Komagataeibacter rhaeticus*

**DOI:** 10.64898/2025.12.15.694344

**Authors:** Jenni Vannas, Amritpal Singh, Bibi Hannikainen, Michelle Alexandrino de Assis, Rodrigo Ledesma-Amaro, Tom Ellis, Rahul Mangayil

**Affiliations:** Department of Bioproducts and Biosystems, Aalto University, Espoo, Finland; Department of Bioengineering, Imperial College London, London, SW7 2AZ, UK; Imperial Centre for Engineering Biology, Imperial College London, London, SW7 2AZ, UK; Bezos Centre for Sustainable Protein, Imperial College London, London W12 0BZ, UK; UKRI Engineering Biology Mission Hub on Microbial Food, Imperial College London, London, W12 0BZ, UK

**Keywords:** Synthetic biology, Microbial chassis engineering, *Komagataeibacter rhaeticus*, Signal peptides, Protein localization, Bacterial nanocellulose

## Abstract

**Background:** Bacterial nanocellulose (BC), produced by *Komagataeibacter* species, is an ideal scaffold for biological Engineered Living Materials (bioELMs) research. Current BC functionalization strategies often rely on secondary microbial hosts or post-production enzyme immobilization, limiting the scalability and modularity required for programmable bioELMs. Establishing a single-chassis system capable of simultaneous biopolymer synthesis and in situ functionalization remains a primary objective in bioELM research. This study addresses the need by benchmarking signal peptide-mediated protein translocation in *K. rhaeticus* iGEM, a model bacterium for BC-based bioELMs, enabling a synthetic biology framework for single-chassis based biomaterial functionalization.

**Results:** Genome-wide analysis confirmed the presence of a complete Sec translocation machinery in *K. rhaeticus*. Through liquid chromatography-tandem mass spectrometry and SignalP 5.0 prediction, native signal peptides were identified and evaluated alongside previously characterized heterologous signal peptides using β-lactamase and mScarlet as cargo proteins. Protein translocation was found to depend on signal peptide identity, cargo type, and expression mode. Fluorescence imaging revealed cytoplasmic, polar, and peripheral localization patterns, confirming functional engagement with the native translocation machinery. A key limitation identified was the retention of recombinant proteins within the periplasm, restricting extracellular availability. Despite this, signal peptide-mediated translocation enabled the incorporation of enzymatic activity into BC during biosynthesis. A post-growth osmotic shock-release strategy increased measurable enzymatic activity by 30%, demonstrating a practical route to overcome this physiological bottleneck while maintaining the biomaterial production capacity.

**Conclusions:** This study benchmarks signal peptide-dependent protein translocation in *K. rhaeticus* and identifies periplasmic retention as a key constraint for extracellular protein release. By linking protein translocation to in situ BC functionalization, this work establishes a synthetic biology framework that supports the development of *K. rhaeticus* as a single-chassis platform for the scalable production of functionalized bioELMs.

## Introduction

Biological engineered living materials (bioELMs) research focuses on integrating living cells with self-assembling biological matrices to create biomaterials with programmable properties [1]. Bacterial nanocellulose (BC), synthesized natively by *Komagataeibacter* spp., offers excellent prospects in bioELM research. Notable features, such as material self-assembly, sustainable production, biodegradability, excellent material properties, and capacity for long-term biocatalyst retention, have made BC an ideal scaffold for bioELM research [2, 3]. A central aspect of bioELM development is biopolymer functionalization. Currently, BC-based bioELM development relies on co-cultivation with an auxiliary protein-secreting microbial chassis or via recombinantly produced enzyme immobilization methods [2]. There remains a need for a single cell chassis capable of both biopolymer synthesis and functionalization, facilitating robust and scalable production of functionalized bioELMs [3].

Recent advances in toolkit development for *Komagataeibacter* spp. have expanded its potential as an engineered chassis [4–6]. For instance, using the Komagataeibacter Tool Kit (KTK), Goosens et al. (2021) engineered heterologous curli (type VIII secretion) system in *K. rhaeticus* iGEM strain and demonstrated extracellular protein assembly and export of β-lactamase [4]. While this demonstrates the feasibility of programming protein export in BC-producing bacteria, the approach relies on complex multiprotein assemblies and is unlikely to be a universal solution. Leveraging native protein translocation pathways offers a potentially simpler and more scalable strategy, yet these remain poorly understood in *Komagataeibacter*.

The Sec translocation pathway is one of the most conserved and extensively characterized protein translocation systems across bacterial lineages and is widely used for directing recombinant proteins across the inner membrane [7–9]. Protein translocation via the Sec translocase occurs through co-translational or post-translational pathways and is primarily governed by signal peptide’s (SP) biophysical properties [10]. SPs recognized by the Sec system contain distinct motifs, positively charged N-terminus, hydrophobic core, and polar cleavage region, which influences protein translocation and localization [11–13]. Systematic benchmarking of SPs compatible with the native *K. rhaeticus* translocation machinery is needed to better understand and guide engineering strategies for BC functionalization.

In this study, candidate SPs native to *K. rhaeticus* were identified using LC–MS/MS-based proteomic analysis combined with bioinformatic prediction. To assess functionality, native and heterologous SPs were fused to two functionally distinct cargos, β-lactamase and mScarlet fluorescent protein. Protein translocation was evaluated under constitutive and inducible expression systems in *E. coli* and *K. rhaeticus*. Translocation dynamics were further examined using time-resolved measurements, and scanning electron microscopy (SEM) was employed to evaluate cell morphology for signs of cellular stress during SP-mScarlet expression. Finally, *K. rhaeticus* expressing SP-β-lactamase fusions were used to generate functionalized BC in pellicle and spheroid forms, with enzyme availability further enhanced using osmotic shock buffer (OSB) treatment.

## Results

### Genome-based identification of Sec translocase machinery and prediction of candidate SPs in *K. rhaeticus*

Genome analysis of *K. rhaeticus* (GCA_900086575.1) confirmed the presence of a complete Sec translocase machinery, including *secA* (2,820,728–2,823,469 bp), *secB* (816,087–816,608 bp), *ftsY* (608,738–609,742 bp), *ffh* (2,305,969–2,307,363 bp), *ffs* (4.5S RNA; 2,502,581–2,502,775 bp), *secY* (2,432,778–2,434,127 bp), *secE* (2,905,499–2,905,774 bp), and *secG* (2,090,741–2,091,046 bp). Presence of these core components indicates *K. rhaeticus* to possess the canonical machinery required for Sec-dependent protein translocation.

To identify native candidate SPs, proteins identified by LC-MS/MS-based proteomic analysis (Table S1) were analyzed using pBLAST. Candidate SPs sequences and their associated translocation pathways, predicted from SignalP 5.0 analysis, were subsequently selected for downstream functional evaluation in *K. rhaeticus* (Table S2).

### Functional evaluation of candidate SPs in *E. coli*

To functionally evaluate the candidate SPs prior to testing in *K. rhaeticus*, each predicted SPs was fused to β-lactamase and constitutively expressed in *E. coli*. β-Lactamase fusions with well-characterized heterologous SPs were included for benchmarking (Table S3).

Nitrocefin assays performed on culture supernatants revealed differences in apparent extracellular β-lactamase activity across SP variants (Figure 1). Among the tested constructs, SP4 showed the highest activity, while YwmC and DsbA were the strongest heterologous benchmarks. Several native *K. rhaeticus* SPs, including SP4, SP9, and SP2, showed comparable activity levels (3.46 ×10^-4^ AU min^-1^, 1.80 ×10^-4^ AU min^-1^, and 1.29 ×10^-4^ AU min^-1^, respectively). However, as *E. coli* does not possess a dedicated system for extracellular release of periplasmically translocated proteins, the measured supernatant activity likely reflects a combination of periplasmic accumulation and passive release. This is supported by the absence of extracellular signals from cytoplasmic β-lactamase and GFP controls, indicating negligible cell lysis.

**Figure 1.**
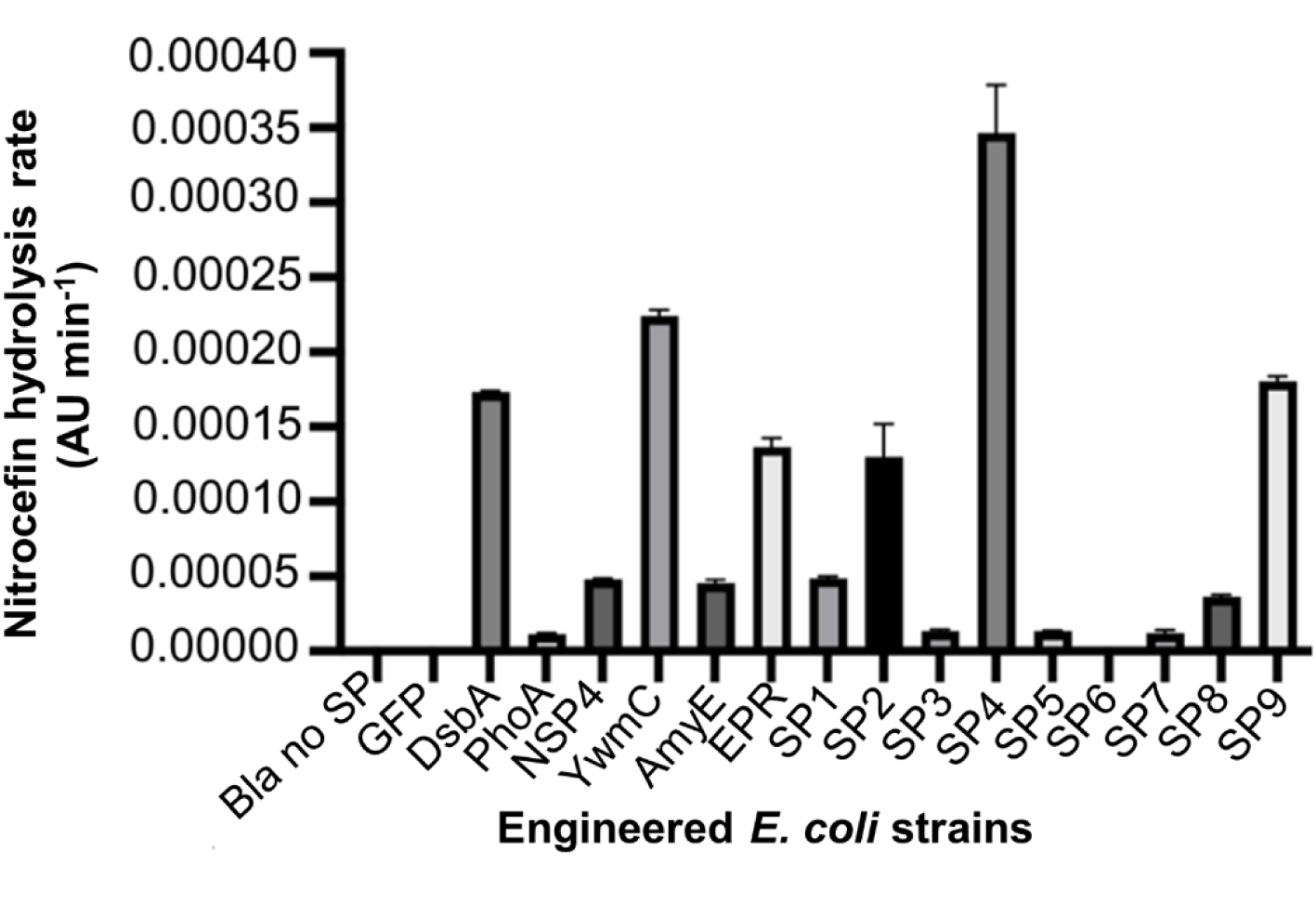
Functional evaluation of candidate SPs in *E. coli*. The graph presents the nitrocefin hydrolysis rate measured from culture supernatants of *E. coli* strains expressing SP-β-lactamase fusions. Control strains included cells expressing β-lactamase devoid of N-terminal SP (Bla No SP) and a green fluorescent protein expression control (GFP). The nitrocefin hydrolysis activity was measured colorimetrically at absorbance of wavelength 480 nm.

### Functional evaluation of candidate SPs in *K. rhaeticus*

The SP- β-lactamase constructs evaluated in *E. coli* were next tested in *K. rhaeticus* using nitrocefin assay (Figure 2). Strains containing PhoA, AmyE, SP1, SP8, and SP10 could not be transformed into *K. rhaeticus* and therefore excluded from further analysis. As expected, a control strain expressing cytoplasmic β-lactamase (Bla no SP) showed negligible activity.

**Figure 2.**
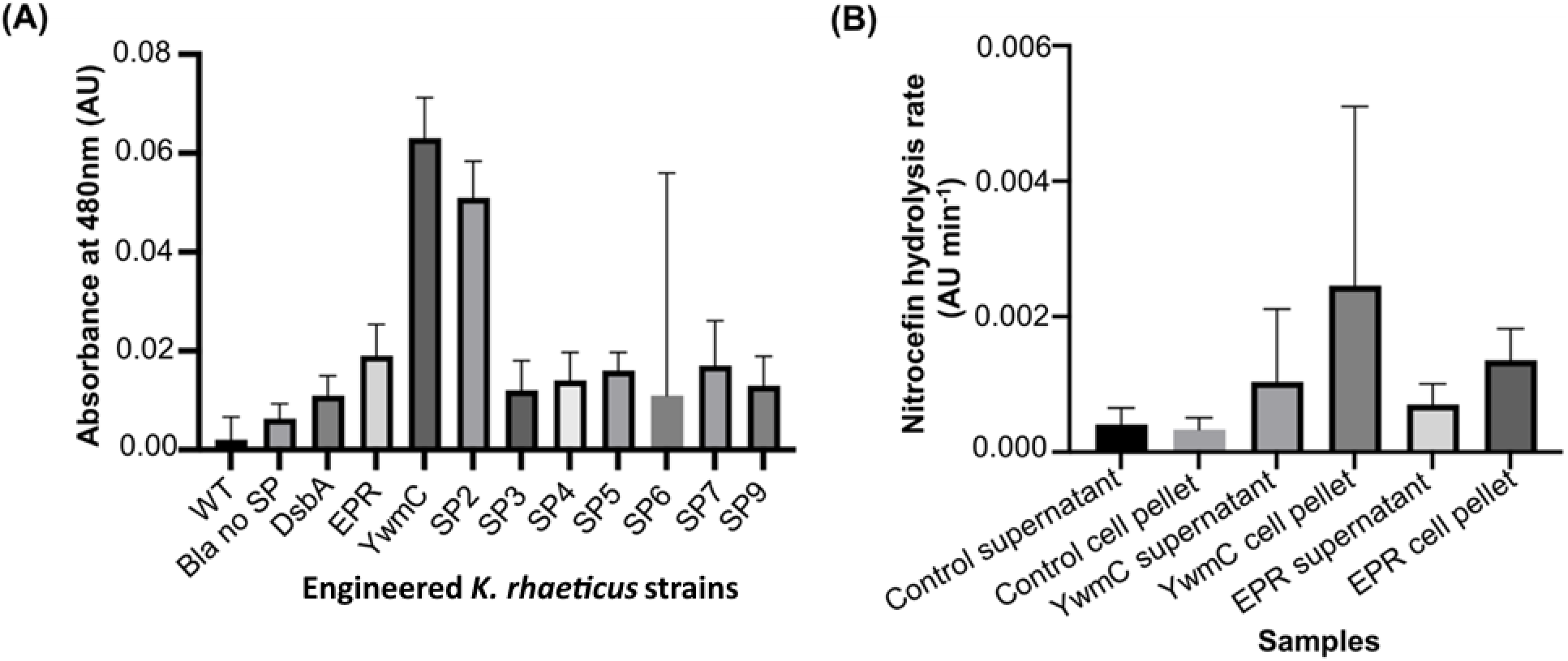
Functional evaluation of candidate SPs in *K. rhaeticus*. (A) Endpoint absorbance at 480 nm measured from cell pellet fractions after 48 h incubation with nitrocefin for strains expressing SP–β-lactamase fusions under a medium-strength promoter (n=3). Wild type (WT) and cytoplasmic β-lactamase (Bla No SP) expressing strains were included as controls. (B) Nitrocefin hydrolysis rates measured from cell pellet and supernatant fractions of cultures expressing YwmC-and EPR-β-lactamase under high-strength promoter, with the cytoplasmic β-lactamase strain as control. Reaction rates are expressed as the change in absorbance at 480nm over time (AU min^-1^). The data points represent the mean value and standard deviations from triplicate samples (n=3).

Analysis of culture supernatants from cells expressing SP-β-lactamase fusions under a medium-strength promoter revealed no detectable activity relative to the control (Figure S1). Consequently, β-lactamase activity was assessed in the cell-associated fractions, where activity was observed for strains expressing EPR-, YwmC-, SP2-, and SP7-β-lactamase, indicating protein expression and translocation into the periplasm (Figure 2A).

To assess whether increased expression could improve extracellular protein availability, the cytoplasmic β-lactamase control and top-performing SP-β-lactamase fusions (Figure 2A) were assembled under a stronger J23104 promoter (Figure 2B). Although SP2 exhibited the second highest activity when expressed under medium strength promoter, variability in culture initiation and expression led to its exclusion, and YwmC- and EPR–β-lactamase fusions were selected for further evaluation. Under these conditions, β-lactamase activity in the culture supernatant reached quantifiable levels (Figure 2B). While the cytoplasmic control showed minimal hydrolysis (3.0 ×10^-4^ AU min^-1^), the supernatant fractions from YwmC-β-lactamase expressing cells exhibited higher hydrolysis rates (1.30 ×10^-3^ AU min^-1^). Notably, cell-associated fractions displayed approximately twofold higher activity relative to the control (Figure 2B). However, as these measurements do not resolve whether the observed activity arises from SP-mediated translocation and extracellular release, or due to cell lysis, subsequent experiments were conducted using SP-mScarlet fusions.

### Evaluation of cargo-dependent protein translocation using constitutive mScarlet expression

To assess SP-dependent protein translocation and evaluate cargo-dependent behavior, SP-mediated translocation of mScarlet fluorescent protein was examined. SP-mScarlet constructs (DsbA, MalE, OmpA, PelB, SP2, SP4, SP10, and YwmC) were assembled under the high strength J23104 promoter in *E. coli*, with cytoplasmic mScarlet serving as a control. Cloning of OmpA- and YwmC-mScarlet fusions under the strong promoter frequently resulted in insertions or deletions within the transcriptional unit (Figures S3 and S4), consistent with toxicity associated with high-level expression. While an intact YwmC-mScarlet construct was obtained using a weaker pLac promoter, OmpA-mScarlet consistently accumulated mutations across promoter variants and was therefore not pursued further under constitutive conditions.

Initial screening in *E. coli* showed growth defects (Figure S4A), with most strains exhibiting growth decline after 24 h, accompanied by reduced intra- and extracellular fluorescence levels (Figures S5A-B and S6A-C), indicating a high metabolic burden associated with SP–mScarlet expression.

Subsequent evaluation in *K. rhaeticus* showed more pronounced growth inhibition compared to *E. coli* (Figure 3A). Only DsbA-, SP2-, SP4-, SP10-, and YwmC-mScarlet strains exhibited measurable fluorescence signals (Figure 3B-C). Culture supernatant fluorescence was generally low and variable across strains, including the cytoplasmic control (Figure 3B, Figure S5). In the case of DsbA-mScarlet, a late increase in fluorescence from culture supernatants coincided with reduced cell growth and increased extracellular signal (Figure 3B-C) suggesting protein release due to compromised cell integrity. In contrast, SP10-mScarlet displayed a delayed fluorescence peak followed by a slower decline, suggesting that the expression of the SP10-mScarlet fusion imposed a reduced cellular burden compared to the heterologous DsbA fusion.

**Figure 3.**
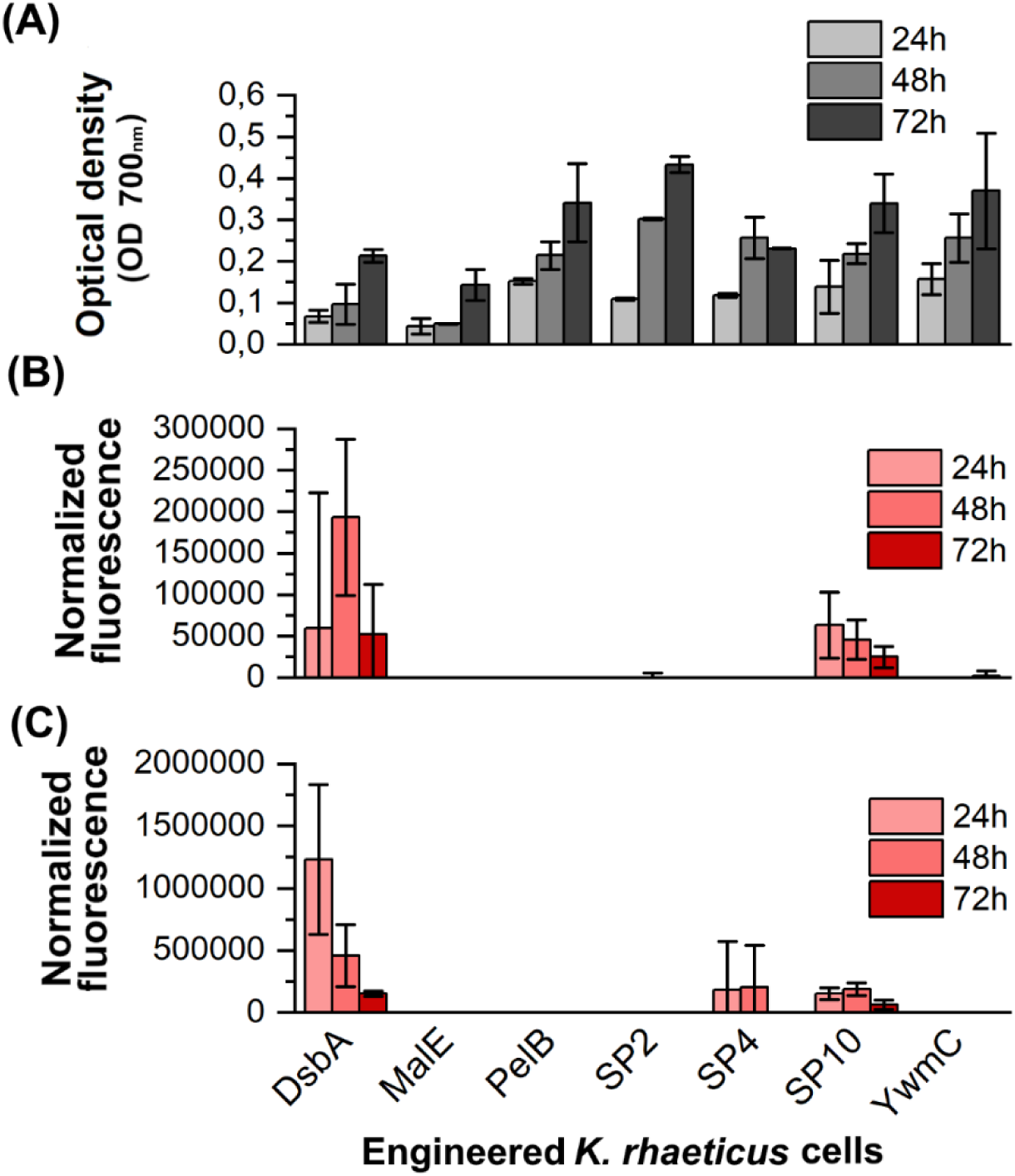
Constitutive expression of SP-mScarlet fusions in *K. rhaeticus.* (A) Cell growth, and normalized fluorescence data from (B) culture supernatant and (C) cell-associated fractions of engineered *K. rhaeticus* cells expressing SP-mScarlet fusions after 24, 48, and 72 h of incubation, respectively. The cytoplasmic mScarlet (-) control showed restricted growth in MA/9 media and was therefore not included. The presented data represents mean values and standard deviations from two biological replicates, each with three technical replicates (n=6).

### SP-mediated mScarlet translocation under regulated expression

To mitigate the physiological burden observed under constitutive expression, SP-mScarlet fusions and the cytoplasmic control were assembled under LuxR/pLux inducible system. Although OmpA-mScarlet could not be constructed under constitutive system, it was successfully assembled in inducible format.

Initial screening in *E. coli* demonstrated improved growth and increased fluorescence signals in culture supernatants (Figure S4D-F and Figure S6). While inducible expression did not fully eliminate growth restriction, it improved fluorescence signals from culture supernatant, allowing subsequent evaluation in *K. rhaeticus*.

In *K. rhaeticus*, inducible expression alleviated the growth inhibition (Figure 4A, Figure S7A-B). Fluorescence signals were detected from the culture supernatants of all engineered strains (Figures S7 and S8), with signal intensities following the order, OmpA > SP4 > YwmC > DsbA > MalE > SP2 > PelB > SP10 (Figure 4B). Several constructs (OmpA, SP4, YwmC, DsbA, SP2) exhibited early peaks in extracellular fluorescence followed by stabilization, whereas MalE-, PelB-, and SP10-mScarlet displayed more gradual accumulation. Cell-associated fluorescence profiles largely mirrored the trends observed in the supernatant (Figure 4C), indicating SP-dependent differences in protein translocation and localization.

**Figure 4.**
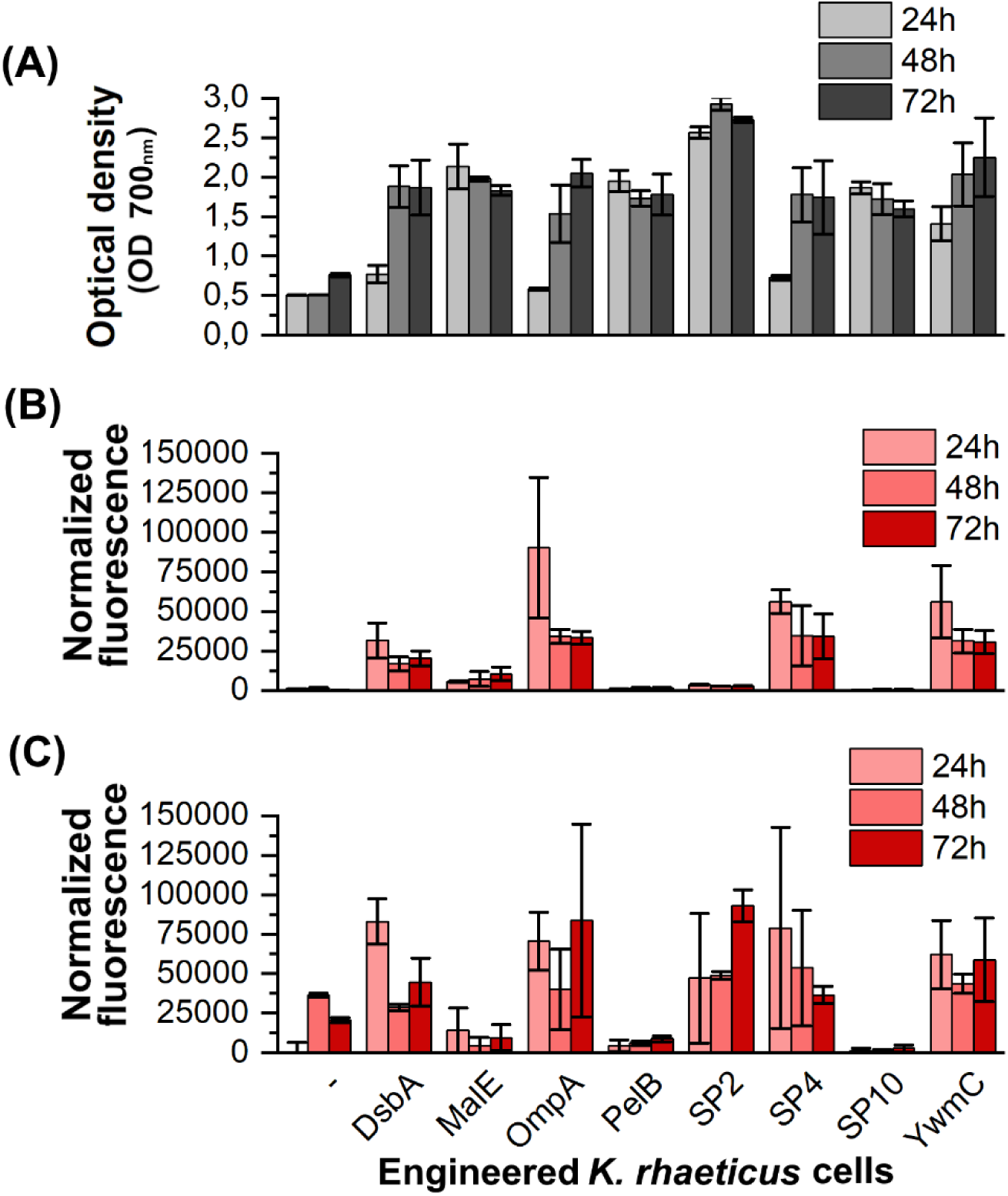
Inducible expression of SP-mScarlet fusions in *K. rhaeticus.* (A) Cell growth, and normalized fluorescence data from (B) culture supernatant and (C) cell-associated fractions of cytoplasmic mScarlet (-) and strains expressing SP-mScarlet fusions after 24, 48, and 72 h, respectively. The data represent mean values and standard deviations from two biological replicates, each with three technical replicates (n=6).

Despite measurable extracellular signals, declines in fluorescence were observed over time. Medium acidification was assessed as a potential factor affecting protein stability. However, HPLC analysis detected low concentrations of metabolic byproducts, such as acetic acid and gluconic acid (Table S4), suggesting that pH effects may not fully account for the observed decrease. Additionally, impaired BC production was observed in SP-mScarlet expressing strains (Figure S9), suggesting that high-level expression and SP-mediated translocation of a rapidly folding cytoplasmic protein impose host-specific physiological constraints, which are potentially associated with periplasmic protein accumulation due to limited release.

### Protein translocation dynamics and morphological analysis of SP-mScarlet expressing *K. rhaeticus*

SP-mScarlet expressing *K. rhaeticus* cells were used to further investigate protein translocation under inducible conditions. To better resolve the decline in apparent extracellular fluorescence between 24 h and 48 h (Figure 4B), short-term fluorescence profiling was conducted over a 10 h period for the best-performing constructs, OmpA- and SP4-mScarlet, alongside the cytoplasmic control (Figure 5).

**Figure 5.**
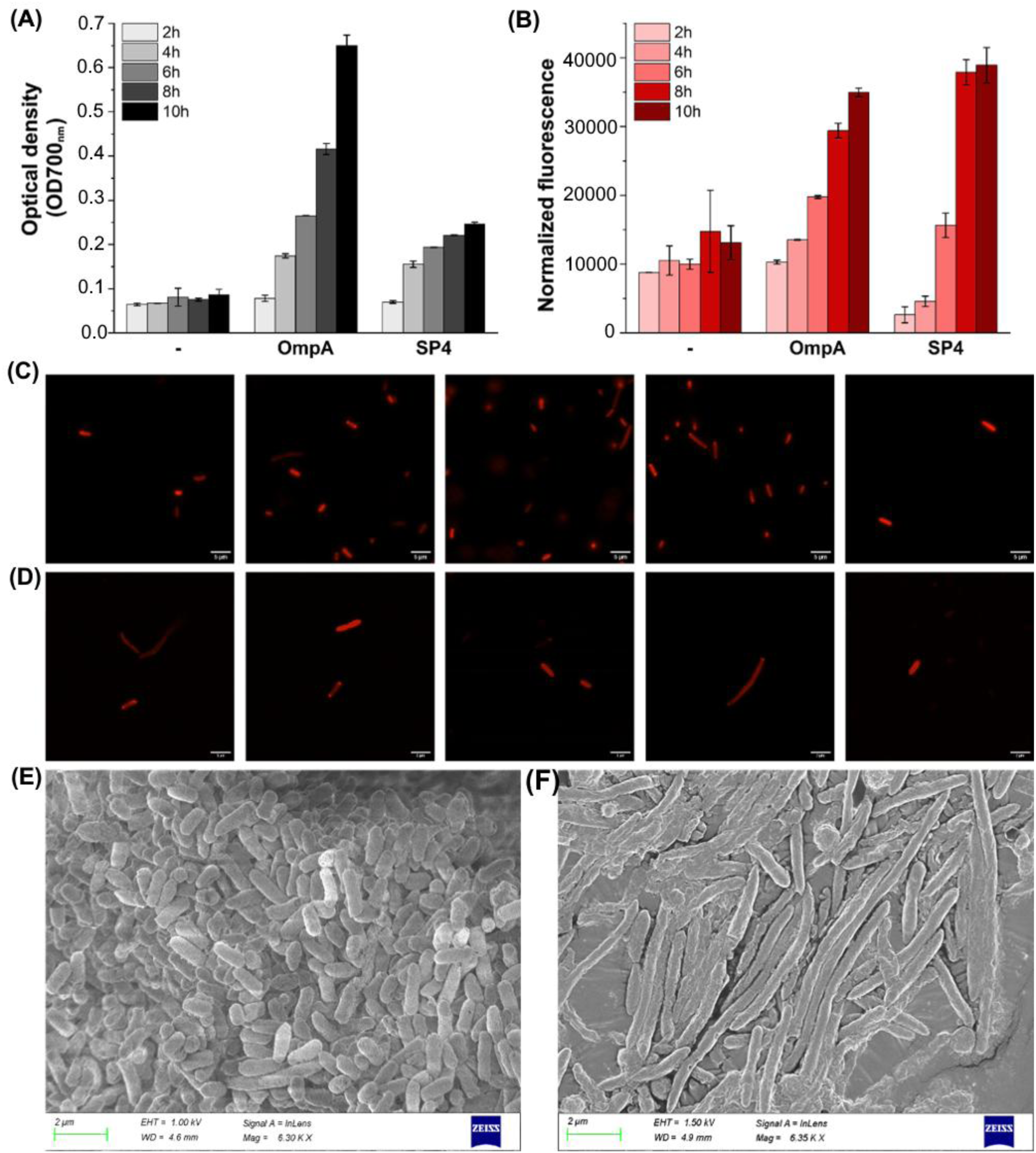
Protein translocation dynamics and morphological analysis of SP-mScarlet expressing *K. rhaeticus* strains. (A) Cell growth and (B) normalized fluorescence signals measured over 2–10 h post-induction. (C-D) Fluorescence microscopy images of OmpA- and SP4–mScarlet-expressing cells, respectively, showing representative localization patterns at indicated time points. (E-F) SEM images of OmpA-mScarlet-expressing cells (E) and cytoplasmic mScarlet control (F) at 10 h post-induction.

Cell growth remained stable for all strains during this interval, with SP-mScarlet expressing cells exhibiting higher overall growth than the cytoplasmic control (Figure 5A). Although fluorescence signals were also observed from the cytoplasmic control culture supernatants, the limited cell growth suggests that the observed signal may be due to stress induced cell lysis (Figure 5A). OmpA-mScarlet expressing cells displayed a steady increase in fluorescence over time and SP4-mScarlet cells showed a more rapid rise followed by stabilization (Figure 5B). Notably, normalized fluorescence levels at 10 h (Figure 5B) were comparable to those measured at 48 h (Figure 4B), indicating that maximal mScarlet occurs during early induction phase.

Fluorescence microscopy revealed three distinct localization patterns from SP-mScarlet expressing cells (Figure 5C-D): (i) diffused cytoplasmic accumulation (OmpA, 2-10 h); (ii) polarized fluorescence (OmpA, 4-8 h; SP4, 2-4 h) suggestive of directed transport toward the cell poles; and (iii) peripheral localization outlining the cell envelope (OmpA, 4-8 h; SP4, 4-10 h), consistent with translocation to non-cytoplasmic compartments.

Although culture supernatant fluorescence levels did not differ significantly between OmpA-and SP4-mScarlet expressing cells (p = 0.85; Figure 5B), OmpA-mScarlet expressing strains achieved higher cell density and was therefore selected for SEM analysis. SEM imaging confirmed that engineered *K. rhaeticus* cells have intact cellular morphology, comparable to the WT cells (Figure 5C; Figure S10B). In contrast, cells expressing the cytoplasmic mScarlet were markedly elongated (Figure 5D; Figure S10), consistent with cellular stress and supporting the interpretation that the extracellular fluorescence observed in the control may arise from compromised cell integrity (Figure 5B).

### In situ BC functionalization via SP-mediated protein translocation and controlled release via OSB treatment

Following SP-dependent protein translocation analyses, the platform was evaluated for in situ BC functionalization. β-Lactamase was selected as the functional module due to its assayable activity, and prior validation across multiple SPs. Constructs incorporating both heterologous (YwmC, EPR) and native (SP4, SP5) SPs were included. Although SP4- and SP5-β-lactamase fusions showed limited extracellular activity under non-cellulose-producing conditions, they were retained based on their ability to support BC pellicle formation (Figure S11).

Except for cells expressing EPR-β-lactamase, all engineered *K. rhaeticus* strains exhibited visible nitrocefin hydrolysis (Figure 6A), indicating the presence of enzymatic activity associated with the material during formation. Given earlier observations of substantial cell-associated retention of recombinant proteins, a “grow-and-release” strategy was implemented, involving BC pellicle formation followed by controlled release using osmotic shock buffer (OSB) [14].

**Figure 6.**
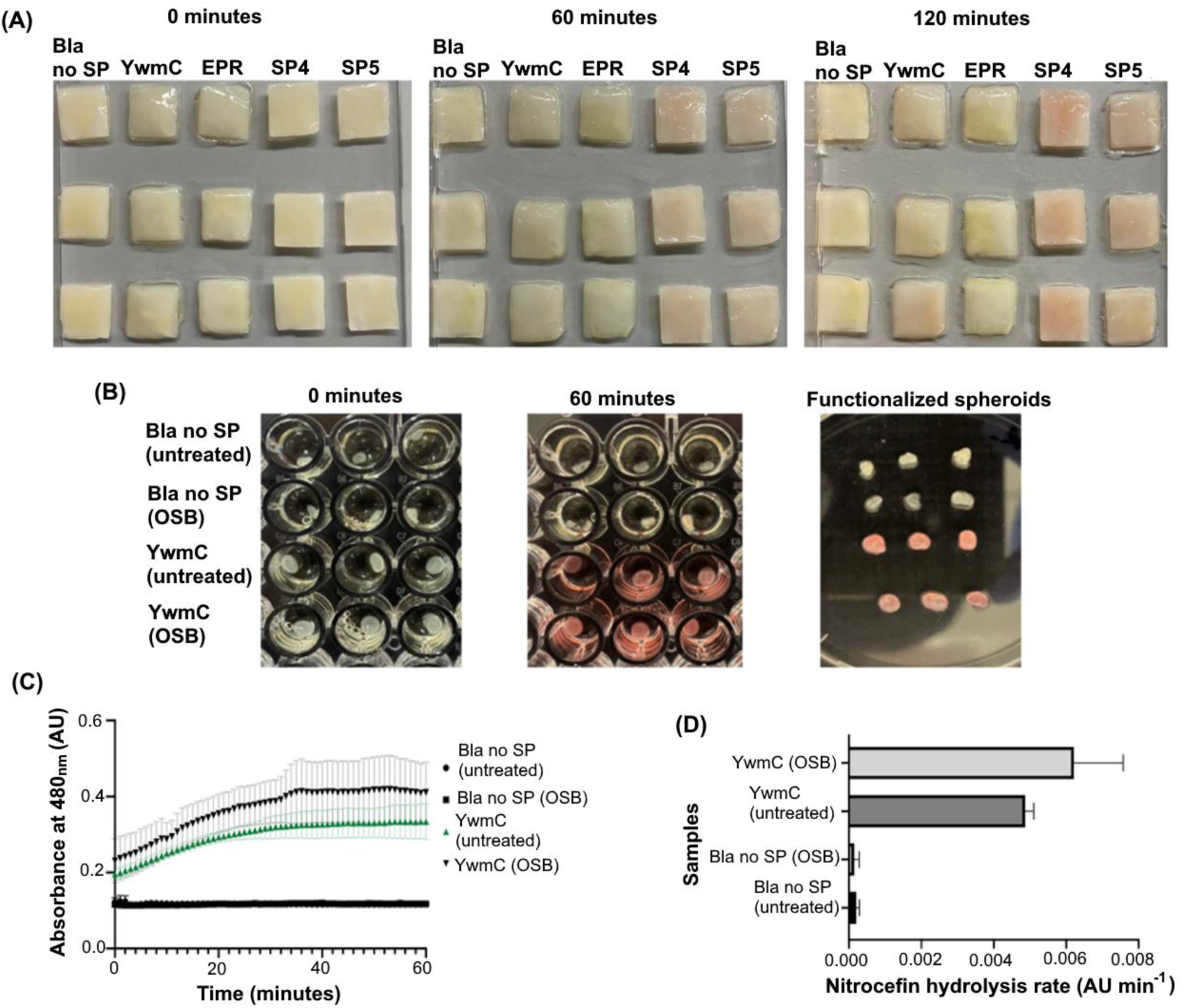
In situ BC functionalization with β-lactamase and controlled release via OSB treatment. (A) BC pellicles grown from engineered *K. rhaeticus* strains expressing YwmC-, EPR-, SP5- and SP4- β-lactamase fusions, along with a cytoplasmic control (Bla no SP). (B) Nitrocefin hydrolysis assay evaluating the effect of OSB treatment on functional activity in BC spheroids synthesized from cytoplasmic control (Bla no SP) and YwmC-β-lactamase. Nitrocefin conversion to its hydrolyzed red form over 2 h provides visual evidence of enzyme activity. BC pellicles and spheroids were washed prior to assay to remove unbound enzyme. (C-D) Quantified hydrolysis rates derived from (B), comparing untreated and OSB-treated spheroids. Reaction rates are expressed as change in absorbance at 480nm over time (AU min^-1^). Data represents standard deviation from triplicate samples (n=3).

To quantitatively assess the effect of OSB treatment, BC production was performed in spheroid format (4-6 mm diameter) [15], facilitating activity measurements in a 96-well plate format (Figure 6B). Spheroid formation was not obtained from EPR-, SP4-, and SP5-β-lactamase expressing strains. However, YwmC-β-lactamase expressing strains and the cytoplasmic control successfully produced spheroids and were further analyzed for controlled release of cell-associated enzyme fractions.

Control spheroids (untreated and OSB-treated) showed no detectable activity, indicating that cytoplasmic β-lactamase is not released under the applied conditions (Figure 6C-D). In contrast, OSB treatment of spheroids derived from YwmC-β-lactamase expressing cells showed increased enzymatic activity from 4.80 ×10^-3^ to 6.21 × 10^-3^ AU min⁻¹, corresponding to an approximately 30% enhancement (Figure 6D).

## Discussion

BC is an attractive scaffold for bioELM research. However, limited knowledge of native protein translocation systems and the lack of genetic tools for recombinant protein secretion in *Komagataeibacter* spp. have necessitated culture optimization and multistep post-production functionalization [3]. Previously Goosens et al. (2021) demonstrated heterologous protein secretion can be achieved in *K. rhaeticus* using engineered systems [4]. However, the relative performance of native and heterologous SPs in this chassis and associated cellular burden remain unclear. This study benchmarks native and heterologous Sec signal peptides to evaluate their capacity to direct protein translocation and localization in *K. rhaeticus*.

Across both *E. coli* and *K. rhaeticus*, apparent extracellular signal levels varied widely among SPs, underscoring the context dependence of SP-mediated protein translocation and localization. Physicochemical principles derived from model organisms suggest that hydrophobic SPs promote translocation efficiency [16–19]. However, this relationship was not strictly conserved in *K. rhaeticus*. Most SPs tested in this study were strongly hydrophobic (GRAVY scores ranging from 0.86 to 1.42, Table S5), yet SP2, despite having a substantially lower GRAVY score (0.221), supported translocation of both cargo proteins. These observations likely reflect a combination of translocation and passive release, as both *E. coli* and *K. rhaeticus* lack efficient mechanisms for extracellular release of periplasmically translocated proteins [20]. Together, these observations indicate that periplasmic retention, rather than translocation efficiency alone, is a major determinant of apparent extracellular protein availability in these systems.

Cargo identity emerged as a major determinant of protein translocation. Despite similar molecular size, SP-β-lactamase constructs remained genetically stable, whereas SP-mScarlet constructs frequently accumulated mutations under constitutive expression (Figure S2-S3). This difference is likely attributable to the expression levels and folding properties of the cargo proteins. Unlike β-lactamase, which is natively secreted and folds in the periplasm, mScarlet is a rapidly folding cytoplasmic protein. Premature folding prior to or misfolding during translocation, as well as overexpression of cytoplasmic proteins, can hinder efficient processing by the host translocation machinery, contributing to cellular stress [21, 22].

Functionally, this bottleneck may extend beyond protein translocation efficiency and influence host physiology. SP-mScarlet-expressing cells exhibited impaired BC production, whereas the cytoplasmic mScarlet and SP-β-lactamase expressing cells retained pellicle formation capacity. This can be interpreted in the context of the cellular organization of BC biosynthesis machinery. The bacterial nanocellulose synthase operon (*bcsABCD*) encodes a trans-envelope complex, with BcsC forming a transmembrane β-barrel channel that mediates glucan-chain export [23–26]. The impaired BC formation in SP-mScarlet expressing cells is likely associated with protein accumulation within the periplasm, where increased protein load may impose spatial or trafficking constraints on the BcsC machinery, which spans the periplasm and outer membrane, thereby impacting glucan-chain export [27, 28]. This suggests a trade-off between recombinant protein load and native biomaterial synthesis, representing a key design constraint for engineering functionalized BC systems.

Although the precise route of protein transport beyond the inner membrane remains unresolved, fluorescence microscopy provides insight into intracellular distribution. The SP-mScarlet expressing *K. rhaeticus* strains exhibited diffused cytoplasmic, polar, and peripheral fluorescence patterns (Figure 5C-D). The diffused cytoplasmic and polar localization patterns are consistent with the ‘*diffusion-and-capture*’ mechanism in rod-shaped bacteria [29], and peripheral signals are consistent with translocation to periplasmic regions following translocation [30].

Maintenance of BC synthesis capacity in SP-β-lactamase expressing *K. rhaeticus* cells facilitated in situ BC functionalization (Fig. 6A-C). Among the successful constructs, native SPs exhibited promising activity under pellicle-forming conditions, suggesting that SP performance is influenced by physiological state (BC forming and planktonic). BC pellicles resemble biofilm-like systems, where protein expression and translocation profiles differ from planktonic conditions [31]. Native SPs may therefore be better adapted to matrix-producing environments, whereas heterologous SPs optimized in planktonic hosts may be less effective under these conditions [32, 33]. However, this observation is based on the tested constructs and does not represent a comparison across SP variants.

A key limitation identified in this study is the retention of recombinant proteins within cell-associated compartments. This was addressed using a grow-and-release strategy, where BC pellicles were subjected to OSB treatments to enhance protein release. As nitrocefin does not cross the inner membrane [34], increased enzymatic activity following OSB treatment indicates the release of non-cytoplasmic β-lactamase. The observed increase in activity (∼30%) confirms that a substantial fraction of recombinant protein remains cell-associated and that this retention can be partially alleviated through post-growth intervention, providing a practical strategy to improve BC functionalization. While the current analysis was limited to constructs that supported robust BC-spheroid formation, these findings demonstrate the feasibility of this approach.

Systematic benchmarking of SPs and the identification of cargo- and host-dependent constraints in *K. rhaeticus* have defined key design principles, specifically regarding the impact of periplasmic retention and expression burden, which will guide the development of a single-chassis platform for BC functionalization. Future efforts in SP engineering and modification of cell envelope interfaces will facilitate more efficient and scalable strategies for in situ BC functionalization across a wide range of biofabrication and material applications.

## Materials and Methods

### Strains and cultivation

Cloning steps, *E. coli* precultures and expression tests were performed with *E. coli* TOP10 (Thermo Fisher Scientific). Recombinant *E. coli* strains were cultured overnight at 30°C (180 rpm) in 5 mL Luria-Bertani (LB) media supplemented with appropriate antibiotics (100 µg mL^-1^ Ampicillin, 34 µg mL^-1^ Chloramphenicol, or 50 µg mL^-1^ Spectinomycin), unless otherwise mentioned. *K. rhaeticus* cells were precultured in 5 mL Hestrin-Schramm (HS) media containing peptone 5 g L^-1^, yeast extract 5 g L^-1^, 2.7 g L^-1^ Na_2_HPO_4_, 1.5 g L^-1^ citric acid, supplemented with 2% glucose, 2% cellulase (Sigma-Aldrich), and appropriate antibiotics (340 µg mL^-1^ Chloramphenicol or 250 µg mL^-1^ Spectinomycin) at 30°C (180 rpm) for 3-5 days as described in Florea et al. [6], unless otherwise mentioned.

### Detection of native *K. rhaeticus* signal peptides

*K. rhaeticus* cells were cultured in 50 mL HS media in 250 mL baffled conical flasks under static conditions at 30°C for 5 days. Culture supernatants were collected and filtered through 0.22 μm pore-size membrane filters. Secreted proteins were concentrated by trichloroacetic acid precipitation by adding 100% TCA solution to the supernatant at a 1:4 (v/v) ratio and incubating overnight at 4 °C. Precipitated proteins were collected by centrifugation at 13,000 rpm for 15 min at 4 °C and washed with ice-cold acetone. After a second centrifugation step, the supernatant was discarded, and the pellet was briefly air-dried. Proteins were subjected to in-solution tryptic digestion by resuspension in 50 mM ammonium bicarbonate (pH 7.8) containing 5 mM dithiothreitol and incubated at 37 °C for 1 h to reduce disulfide bonds. Iodoacetamide was added to a final concentration of 15 mM, and samples were incubated for 30 min at room temperature in the dark for alkylation. The reaction mixture was diluted with three volumes of 50 mM ammonium bicarbonate (pH 7.8), and pH was adjusted where necessary using 50 mM Tris–HCl (pH 8.0). Trypsin Gold (Promega) was added at a 1:20 (w/w) protease-to-protein ratio, and digestion was carried out overnight at 37 °C. The reaction was terminated by adding formic acid to a final concentration of 1% (v/v), and peptide mixtures were analyzed by liquid chromatography-tandem mass spectrometry (LC-MS/MS).

LC–MS/MS analysis was performed by Dr. David Bell at the Proteomics Facility, Imperial College London, who provided peptide identification data. Identified peptides were matched against the SwissProt database and the *K. rhaeticus* genome (GCA_900086575.1) to assign protein identities. The corresponding protein sequences were analyzed using SignalP 5.0 for N-terminal SP prediction, specifically for Gram-negative bacteria [35, 36]. Selected SPs were synthesized as gBlocks (Integrated DNA Technologies) and cloned into KTK D1.1 entry vectors (Table S6) as previously described [4].

### Plasmid transformation

Chemically competent *E. coli* TOP10 (Thermo Fisher Scientific) cells were handled as described in Sambrook and Russell [37]. Colony PCR using appropriate KTK level- primers and whole plasmid sequencing (Eurofins) were used to verify the constructs. *K. rhaeticus* cells were transformed by electroporation as described in Florea et al. [6]. GC prep developed by Blount et al. (2016) [38] was used for colony PCR confirmation using specific KTK primers [4]. The list of KTK parts, plasmids and KTK specific colony PCR primers used in this study are presented in Tables S6-S8, respectively.

### *E. coli* and *K. rhaeticus* expression culture conditions

Expression cultures in MA/9 media were initiated by inoculating pre-cultivated cells to a starting optical density at wavelength 700 nm (OD_700nm_) of 0.05. The MA/9 contained 200 mL L^-1^ MA/9 5x salt stock solution (27.6 g L^-1^ Na_2_HPO_4_ * 2H_2_O, 17 g L^-1^ KH_2_PO_4,_ 5 g L^-1^ NH_4_Cl, 0.04 g L^-1^ nitriloacetic acid, and 5 g L^-1^ NaCl in MQ), 2 mL L^-1^ 1M MgSO_4_, 0.1 mL L^-1^ 1M CaCl_2_, 0.05 mL/ L^-1^ FeCl_3_ (10 mg mL^-1^), and 10 mL L^-1^ casamino acids (20% m/V) in MQ and supplemented with 2% glucose, appropriate antibiotics (Chloramphenicol or Spectinomycin for constitutive and inducible expression, respectively), and 50 µM AHL (N-(β-Ketocaproyl)-L-homoserine lactone) (Sigma-Aldrich) upon start of experiment. Cultures were grown in biological triplicates at 30°C with shaking (180 rpm) for 24, 48, and 72 hours.

### Nitrocefin assay

Nitrocefin (chromogenic β-lactamase substrate) (Merck) was prepared as a 10 mg mL^-1^ stock solution in dimethyl sulfoxide (DMSO) according to the manufacturer’s instructions. Immediately prior to use, the stock was diluted to a working concentration of 100 μg mL^-1^ in nitrocefin assay buffer (50 mM sodium phosphate, 1 mM EDTA, pH 7.4). Overnight cultures of *E. coli* and *K. rhaeticus* were grown and adjusted to OD_600nm_ of 0.6-0.8. For β-lactamase activity measurements, 50 µL of nitrocefin working solution was combined with 50 µL of either cell culture or culture supernatant in individual wells of a flat-bottom 96-well microtiter plate. Plates were incubated at room temperature in a microplate reader, and nitrocefin hydrolysis was monitored by measuring the absorbance at 480 nm at defined time intervals. Assays were conducted for up to 2 h for *E. coli* and up to 48 h for *K. rhaeticus*.

### Nitrocefin assay using BC pellicles

*K. rhaeticus* BC pellicles were produced using a standard static culture protocol in 24-deep-well plates containing 5 mL HS media, with engineered strains grown in the presence of chloramphenicol (340 μg mL^-1^). After incubation for 5 days at 30°C, pellicles were carefully removed and subjected to three sequential washing steps to remove residual medium and loosely associated cells. For each wash, individual pellicles were transferred to 5 mL sterile phosphate-buffered saline (PBS), vortexed at maximum speed for 1 h at room temperature, after which the wash solution was discarded and replaced with fresh PBS. Following the final wash, pellicles were transferred to sterile Petri dishes, and 50 μL of nitrocefin working solution was applied to fully cover the pellicle surface. β-Lactamase activity was assessed qualitatively by monitoring nitrocefin conversion, observed as a yellow-to-red color change on the pellicle surface. The reaction was documented by photography at defined time points over a 2 h period using a standardized imaging setup.

### Nitrocefin assay using *K. rhaeticus* BC spheroids

*K. rhaeticus* spheroids were generated culturing cells at 250 rpm for 3 days without cellulase treatment. Spheroids were harvested and subjected to three consecutive washing steps to remove residual medium components and loosely associated cells. For each wash, spheroids were transferred to 5 mL sterile PBS, vortexed at maximum speed for 1 h at room temperature. The wash solution was then discarded and replaced with fresh PBS for each cycle. Following washing, individual spheroids were transferred into wells of a 96-well microtiter plate. Nitrocefin working solution (50 µL per well; prepared according to the assay method) was added directly to each spheroid-containing well, and β-lactamase activity was monitored by measuring the increase in absorbance at 480 nm over 60 min using a microplate reader.

### OSB treatment of BC spheroids

The OSB buffer contained 200 mM Tris-HCl, 1 mM EDTA, 20% (w/v) sucrose and 500 μg mL^-1^ lysozyme. BC spheroids were prepared, harvested, and washed as described above, and then transferred into OSB diluted to 25% (v/v). Samples were incubated at room temperature for 1 h, after which the nitrocefin assay was performed as described previously.

### Fluorescence analysis

Cell growth (OD_700nm_) and mScarlet fluorescence (ex/em 569/594nm) were measured in triplicates in 96-well plates using Cytation Reader 3 (Biotek). Cultures (5 mL) were centrifuged at 3900 rpm for 10 min to separate supernatant and cell fractions. Supernatants were filtered through 0.2 µm PTFE filters (Captiva Econo PTFE, Agilent Technologies) using 1 mL Injekt-F syringes (Braun). Cell pellets were resuspended in 1 mL of 1X sterile PBS. MA/9 medium and PBS were respectively used as blanks in supernatant and cell-associated fraction measurements. Fluorescence values were normalized to cell density and reported as RFU/OD_700nm_.

Filtered supernatants were further analyzed for pH, HPLC, and SDS–PAGE. Cell pellets were used for SDS–PAGE, fluorescence microscopy, and SEM analysis. Supernatant pH was measured using SevenCompact pH meter with InLab Micro pH probe (Mettler Toledo).

### Analytics

Filtered supernatant samples from recombinant *K. rhaeticus* cultivations were diluted 1:10 and analyzed by HPLC (Ultimate 3000, ThermoFisher Scientific) equipped with UV and refractive index detectors. Separation was performed using a Rezex ROA-Organic Acid H⁺ column (8%, 300 × 7.8 mm; Phenomenex) maintained at 55 °C. The mobile phase consisted of 5 mM H₂SO₄, delivered at a flow rate of 0.5 mL min⁻¹. UV detection was performed at 210 nm.

Fluorescence microscopy was performed using an Axio Observer Z1 microscope (Carl Zeiss, Germany) equipped with a 16×/0.3 objective, a PvCam camera, and a Colibri.2 light source. mScarlet fluorescence was detected using excitation at 594 nm. Z-stack images were processed using Fiji [39].

SEM was conducted using a Zeiss Sigma VP microscope equipped with an InLens detector. Cell pellet samples were fixed overnight in 2.5% (v/v) glutaraldehyde (prepared from a 25% solution; Sigma-Aldrich) at a volume 10–20× that of the pellet. Samples were subsequently dehydrated using graded ethanol washes (30% and 50%), and chemically dried overnight using 1 mL hexamethyldisilazane (HMDS; Sigma-Aldrich). Dried samples were mounted on SEM stubs and sputter-coated with a 4–5 nm Au–Pd layer using a Leica EM ACE600 system.

## Supporting information

Supplementary figures and tables are included in the supporting information file.

## Author Contributions

The manuscript was written through contributions of all authors. All authors have given approval to the final version of the manuscript. ‡These authors contributed equally.

## Acknowledgments

The authors acknowledge Dr David Bell at the proteomics suite at Imperial College London and Sarvesh Kumar at Aalto University for conducting the LC-MS/MS and HPLC analysis, respectively.

