## Supplementary File for "A signal peptide-guided platform for in situ functionalization of bacterial nanocellulose in *Komagataeibacter rhaeticus*"

**Supplementary Figures**

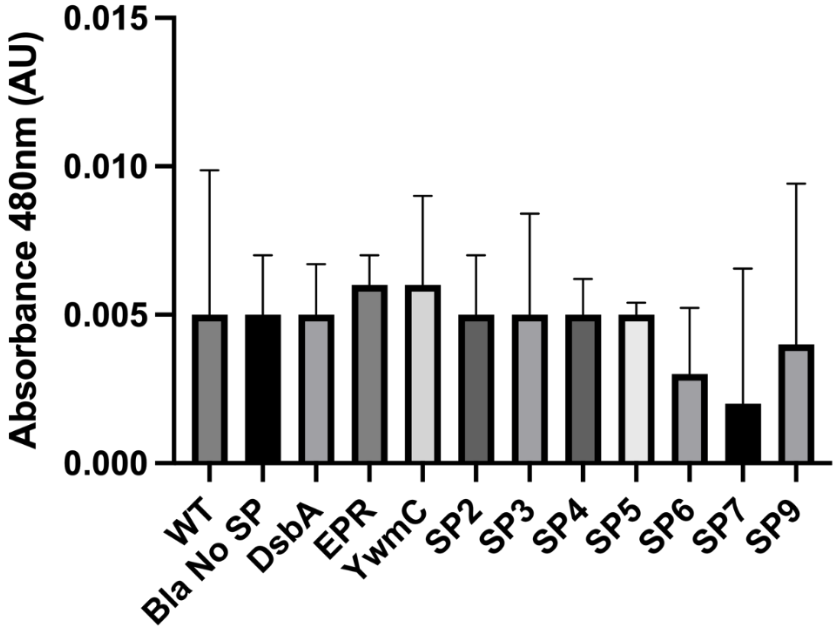


Figure S1. Endpoint nitrocefin assay performed on culture supernatants of engineered *K. rhaeticus* strains and controls (WT, Bla No SP) after 72 h of cultivation.

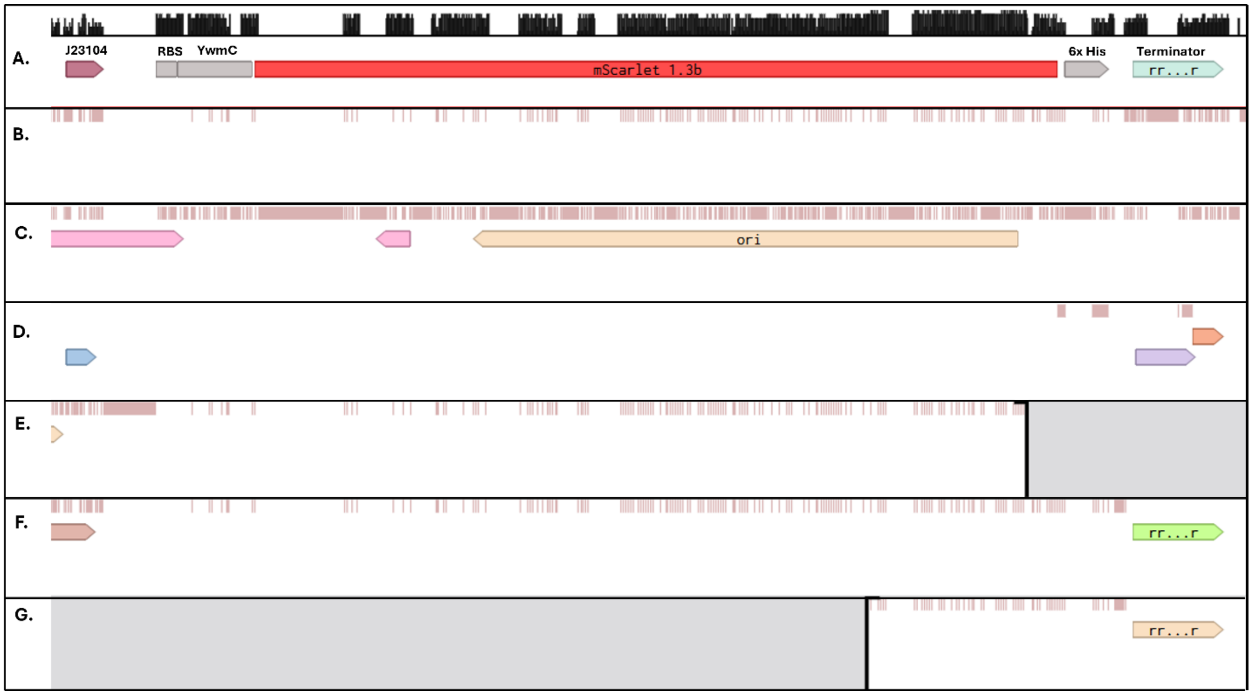


Figure S2. Sequencing results of constitutively expressed YwmC-mScarlet constructs revealing insertions and deletions in the transcriptional unit. Comparison to (A) template sequence revealed that the assemblies carried mutations especially in the (B, E-F) promoter, (B, E-F) mScarlet coding sequence, and (D) C-terminus of the fusion protein and terminators. Alternatively, substitution of the entire transcriptional unit with fragments from the (C) plasmid origin of replication and (G) deletion of majority of the coding sequence were observed.

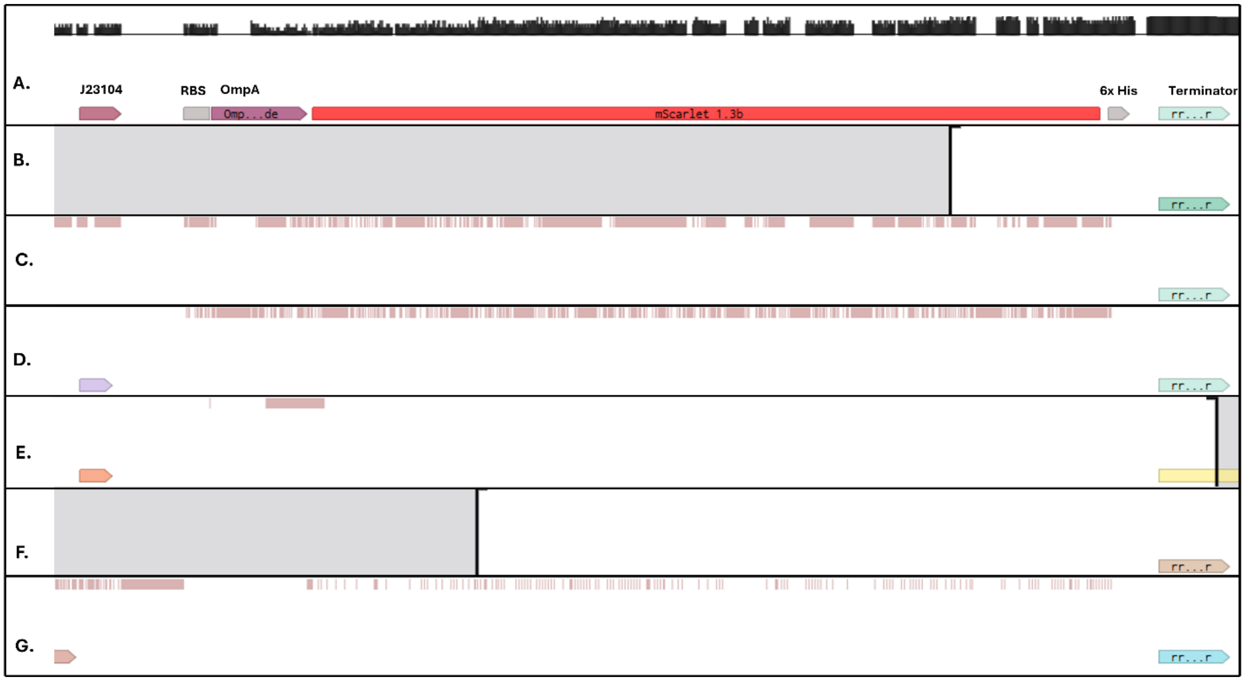


Figure S3. Sequencing results of constitutively expressed OmpA-mScarlet constructs revealing insertions and deletions in the transcriptional unit. Comparison to (A) template sequence revealed that the assemblies carried mutations especially in the (C, G) promoter, (C-D, G) OmpA-mScarlet coding sequence, and (E) OmpA sequence. (B, F) Additionally, deletions of most of the coding sequence were observed.

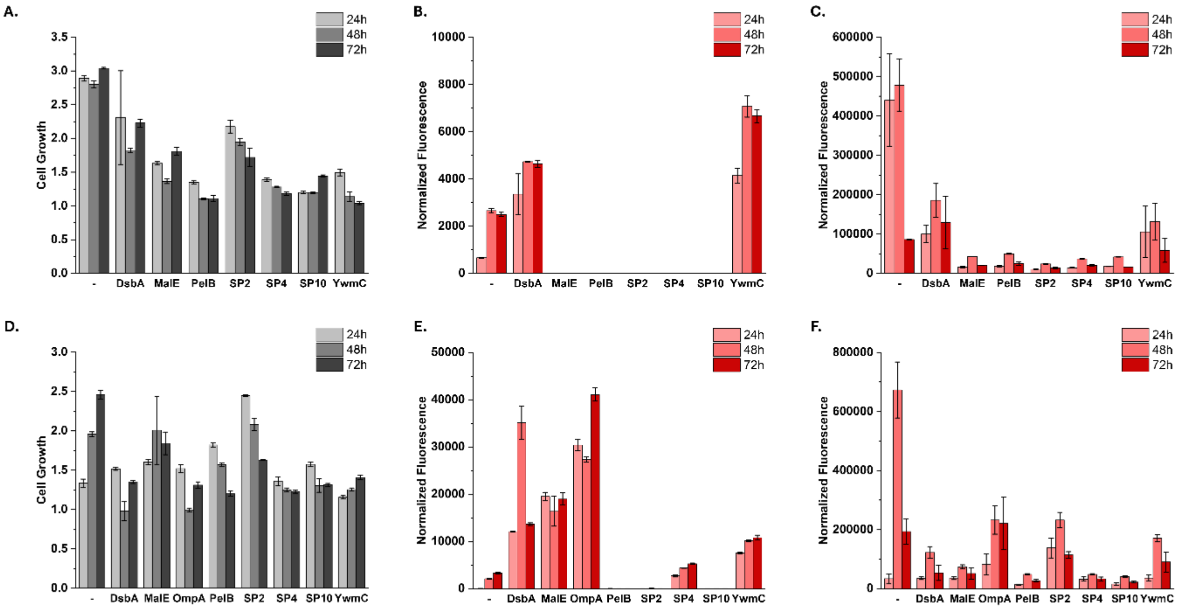


Figure S4. Constitutively expressed SP-mScarlet fusions in *E. coli.* The subfigures present (A) cell growth ($OD_{700nm}$), (B) normalized fluorescence from culture supernatant, and (C) cell-associated fractions of cytoplasmic mScarlet (-) and strains expressing SP-mScarlet fusions after 24, 48, and 72 hours, respectively. (D-F) presents the corresponding data for inducibly expressed SP-mScarlet fusions. The presented data represents mean values and standard deviations from two biological replicates, each with three technical replicates (n=6).

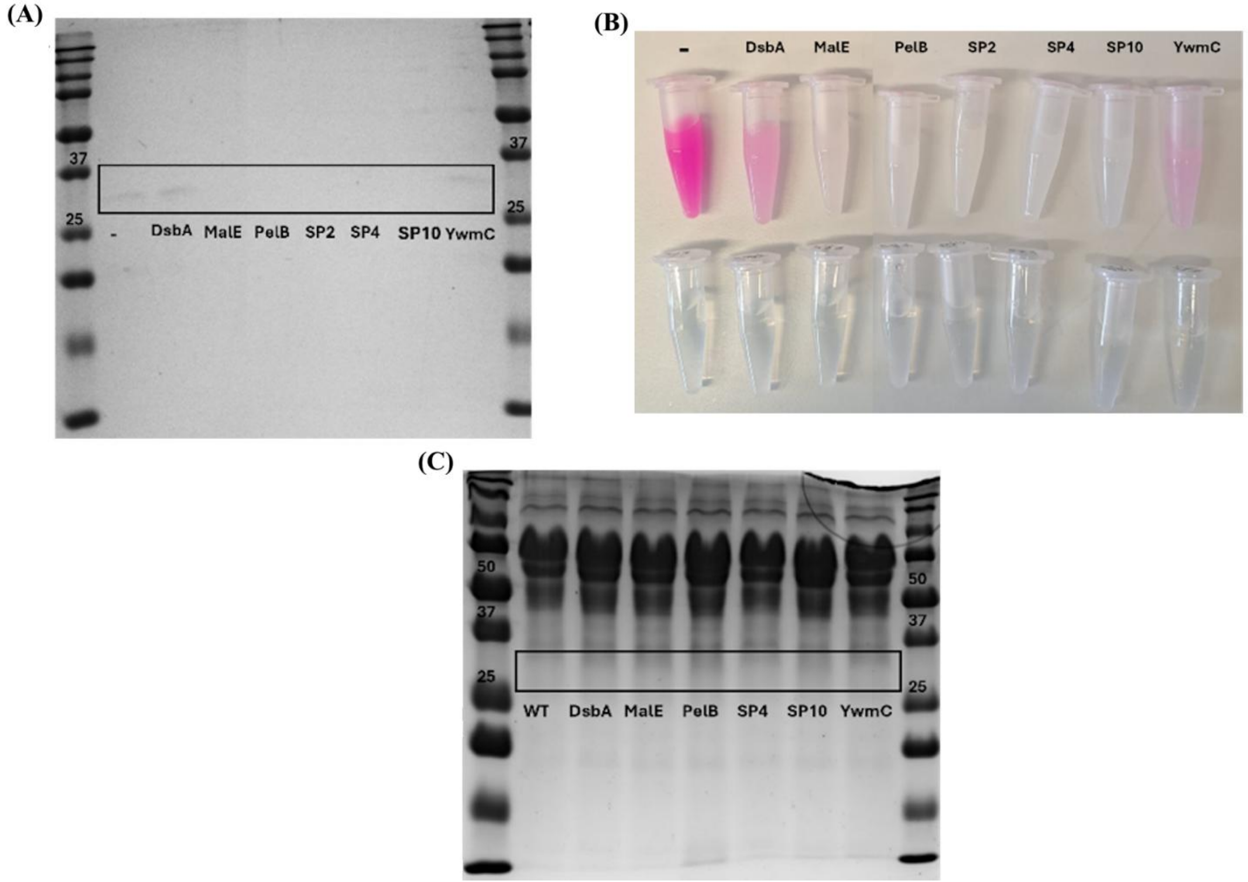


Figure S5. Constitutive expression of cytoplasmic mScarlet (-) and SP-mScarlet fusions in *E. coli*. (A) SDS-PAGE analysis of culture supernatants assessing the presence of mScarlet (26.4 kDa). (B) Visual appearances of recombinant E. coli cells after 24 h of constitutive expression. Pellets (intracellular fluorescence) and supernatants (extracellular fluorescence) are presented on the top and bottom rows, respectively. (C) Constitutive expression of cytoplasmic mScarlet (-) and SP-mScarlet fusions in *K. rhaeticus*. SDS-PAGE showing no detectable mScarlet in culture supernatants (26.4 kDa). Exogenous cellulase can be observed as large bands around 68 kDa.

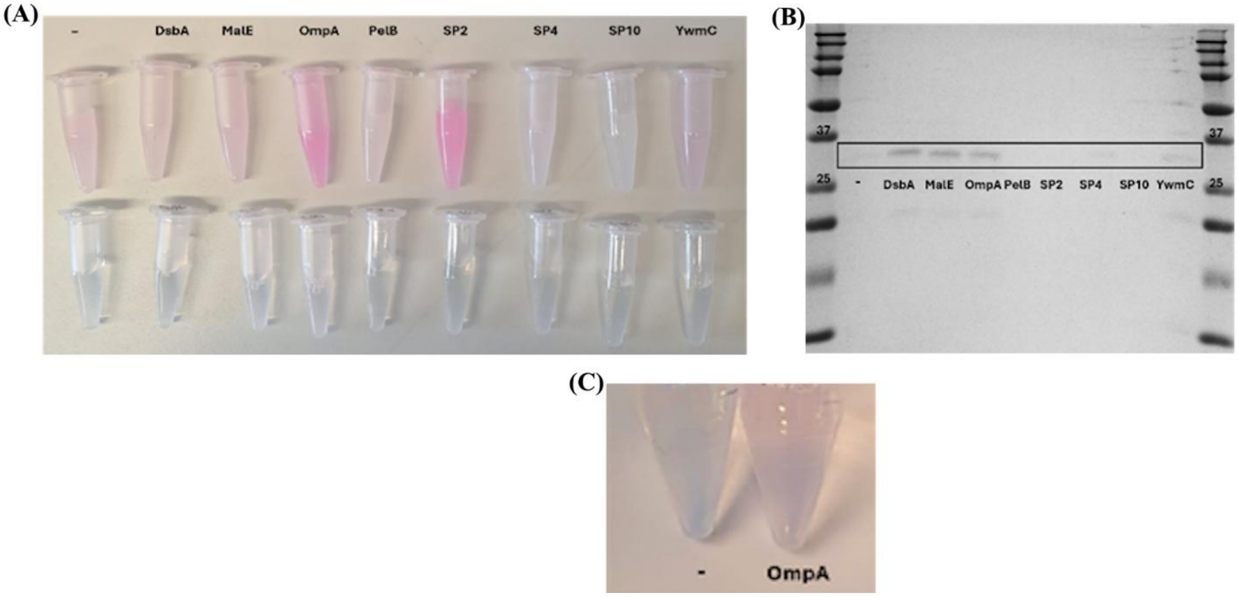


Figure S6. Inducible expression of cytoplasmic mScarlet (-) and SP-mScarlet fusions in *E. coli*. (A) Visual appearance of inducible expression strains at 24 h. Pellets on top row and supernatants bottom row. (B) SDS-PAGE analysis of culture supernatants for detection of mScarlet (26.4 kDa). (C) A closer look at supernatant color differences between cytoplasmic mScarlet (-) and OmpA-mScarlet.

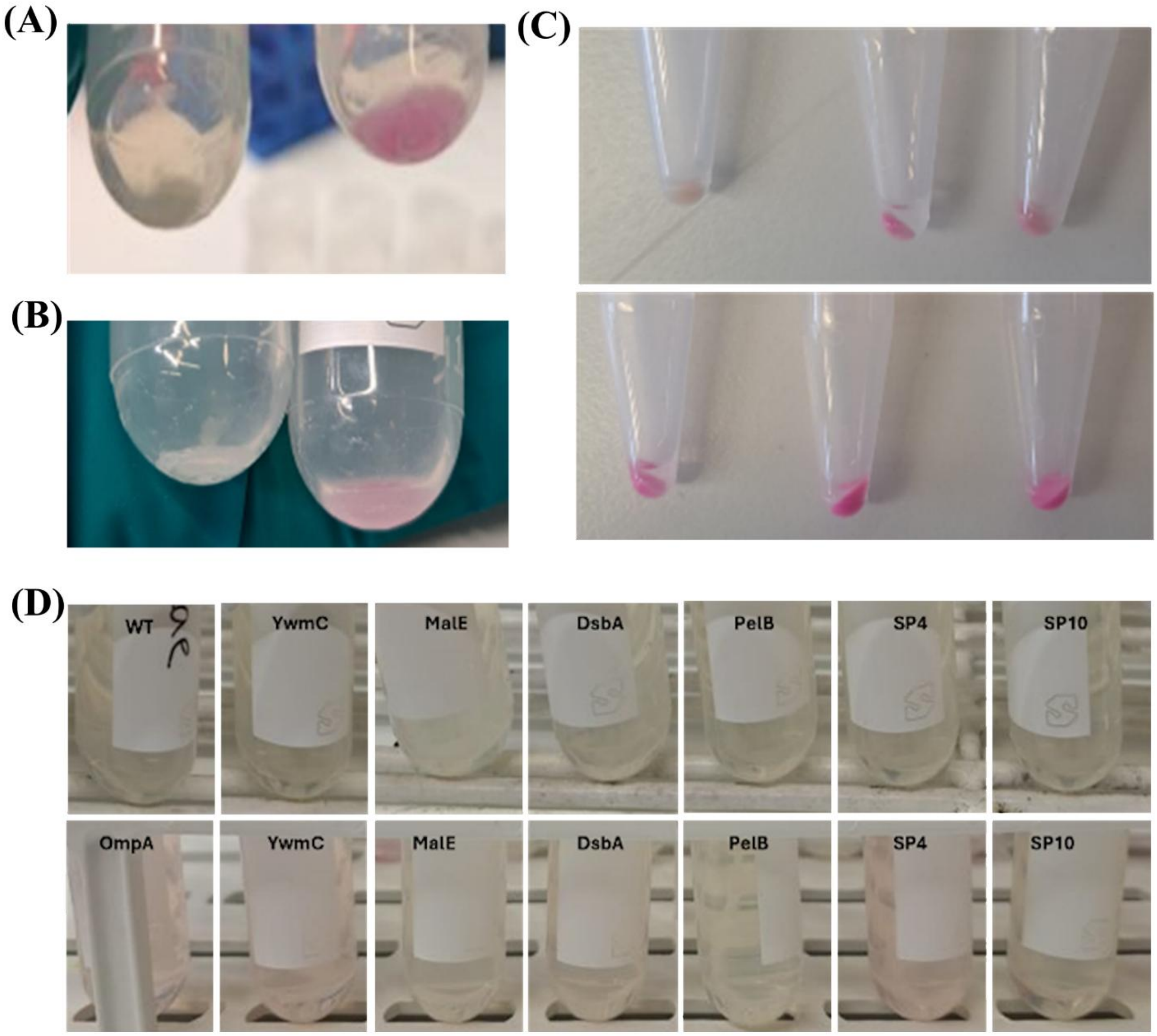


Figure S7. Visible differences in the constitutive and inducible expression strains. Pellet size and color difference between constitutive (left) and inducible (right) (A) SP4-mScarlet and (B) YwmC-mScarlet after 48 h of induction. (C) Pellet fractions of inducible cytoplasmic mScarlet (top row) and OmpA-mScarlet (bottom row) after 24 h, 48 h, and 72 h of induction, from left to right respectively. (D) Filtered supernatants of cells expressing wild-type (WT), and constitutive (top row) and inducible (bottom row) signal peptide-mScarlet fusions.

(A)

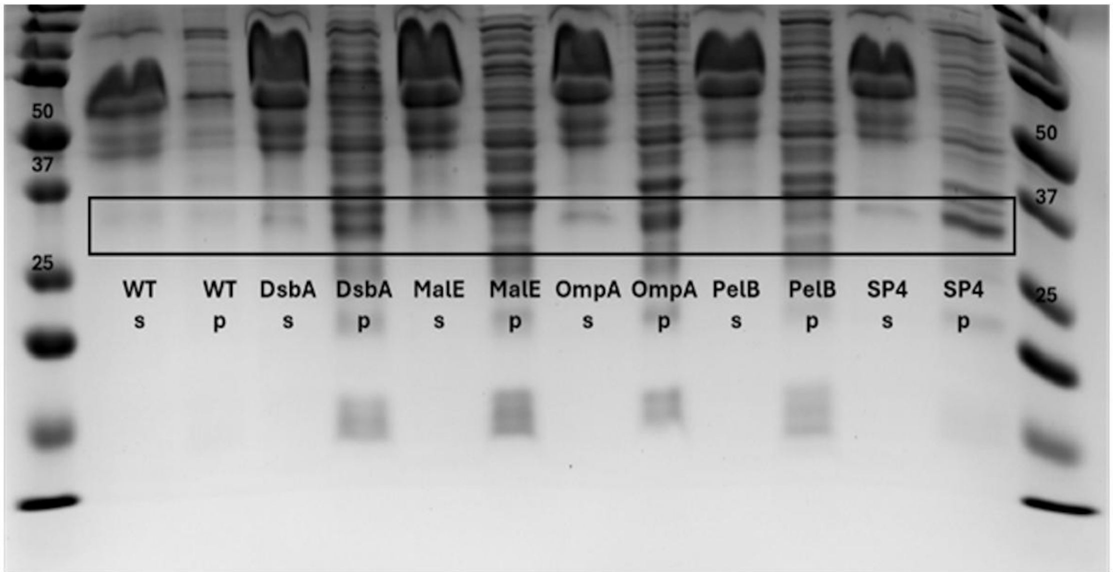


(B)

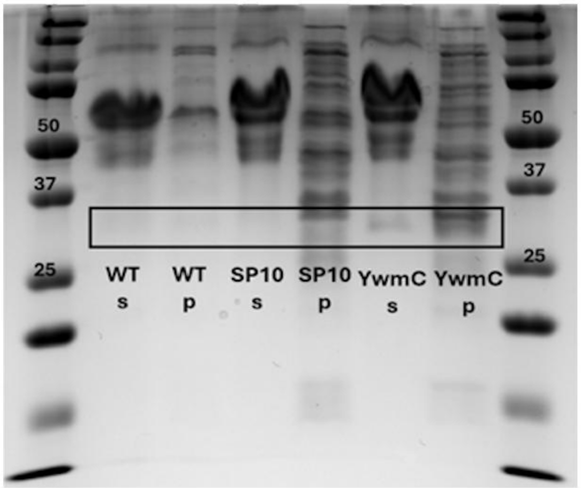


Figure S8. SDS–PAGE analysis of inducible SP-mScarlet expression in *K. rhaeticus*. (A-B) Comparison of culture supernatant (s) and cell pellet (p) fractions for strains expressing SP-mScarlet fusions. Bands corresponding to SP-mScarlet (29.7–31.6 kDa; Table S5) and cytoplasmic mScarlet (26.4 kDa) are highlighted. Strong signals are primarily observed in pellet fractions, indicating intracellular or cell-associated protein accumulation, while only faint bands are detected in supernatant fractions. A prominent band at ~68 kDa corresponds to supplemented cellulase present in the medium.

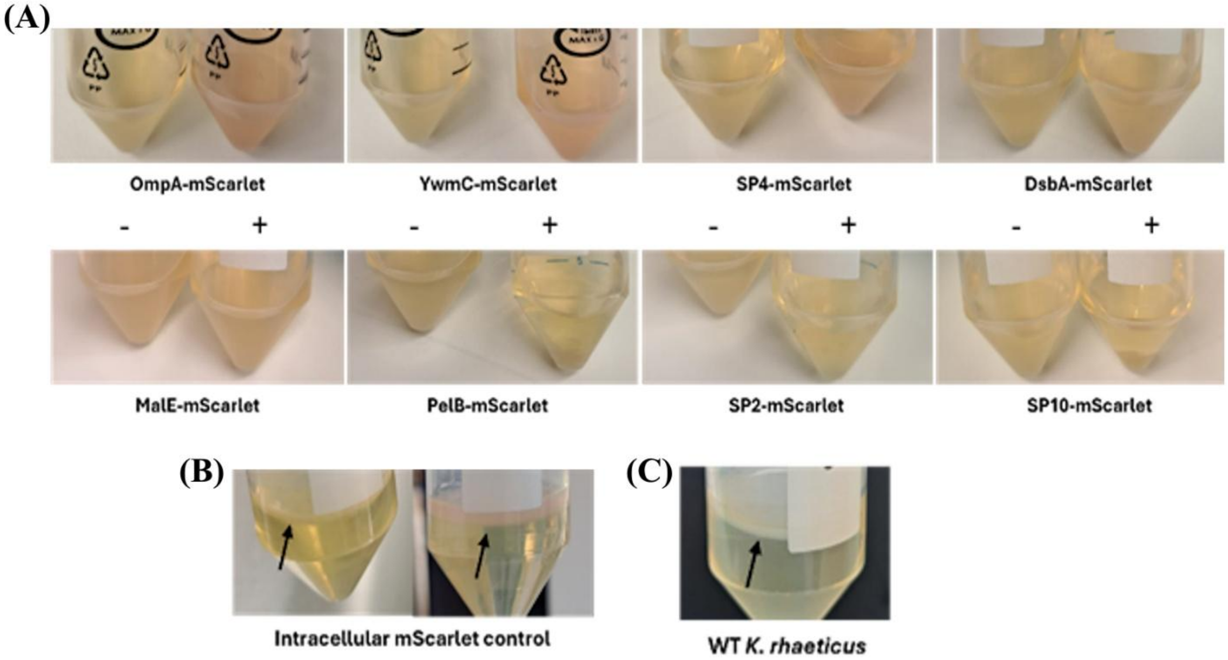


Figure S9. Five-day static cultivation of SP-mScarlet expressing *K. rhaeticus* cells. (A) Inducible SP-mScarlet and (B) mScarlet control strains without (-) and with (+) AHL induction did not exhibit a defined cellulose pellicle. (C) Cellulose pellicle of WT *K. rhaeticus* strain.

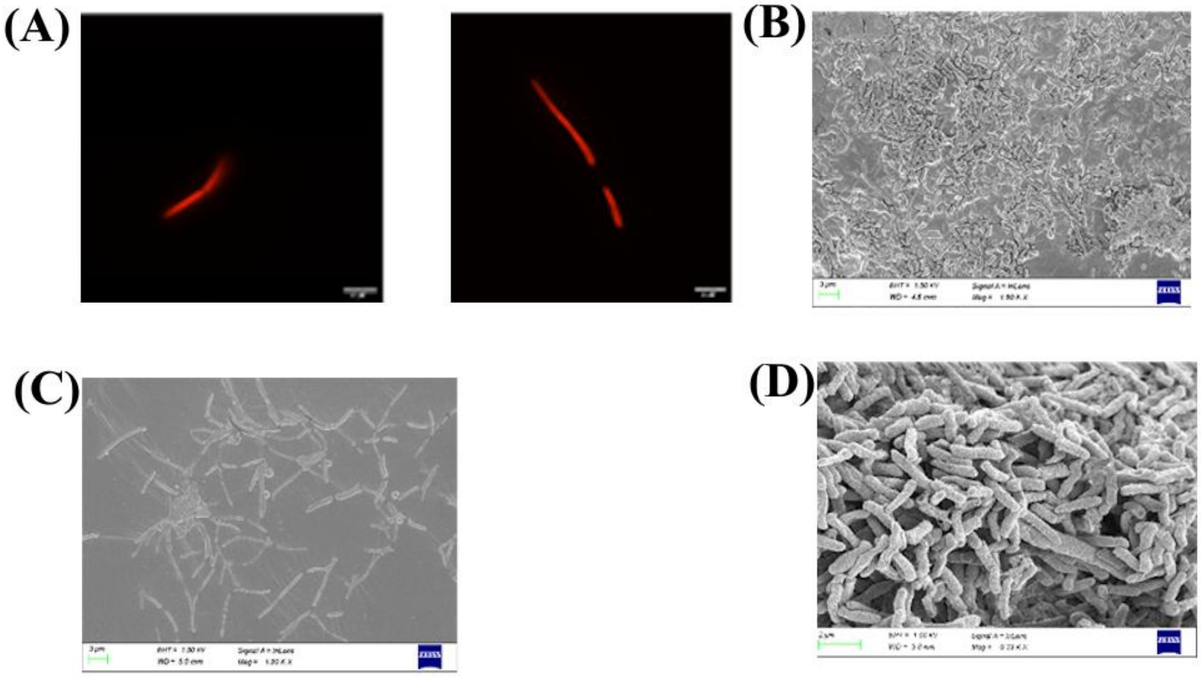

Figure S10. Microscopy analysis of recombinant *K. rhaeticus*. (A) Fluorescence microscopy images of cytoplasmic mScarlet after 6 h and 10 h post-induction shows elongated cell morphologies. Wider scanning electron microscopy (SEM) images of (B) OmpA-mScarlet and (C) cytoplasmic mScarlet support the difference in morphologies observed during fluorescence microscopy. SEM analysis of (D) WT *K. rhaeticus* after 10 h of culturing.

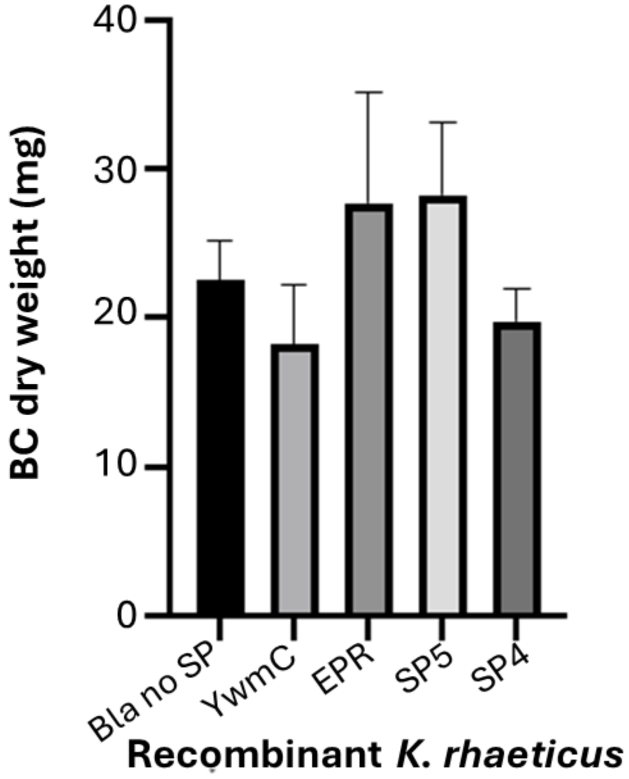


Figure S11. BC production by *K. rhaeticus* cells expressing SP-β-lactamase.

### SUPPLEMENTARY TABLES

Table S1. Proteins identified by LC-MS/MS proteomic analysis.

| **Proteins** | **Cellular Localization** |
|---|---|
| Putative Endoglucanase CMCAX_*(SP6) | Extracellular |
| Toluene transporter subunit (TodC1)_*(SP7) | Periplasmic |
| Hypothetical Protein 1/truncated endoglucanase_*(SP1) | N/A |
| Virginiamycin B lyase_*(SP8) | N/A |
| Hypothetical Protein 2_*(SP2) | N/A |
| Hypothetical Protein 3_*(SP3) | N/A |
| Outer Membrane Protein OmpA_*(SP10) | Extracellular* |
| Periplasmic pH-dependent serine endoprotease DegQ_*(SP9) | Periplasmic |
| Hypothetical Protein 4_*SP(4) | N/A |
| Alcohol Dehydrogenase 15kDa Subunit | N/A |
| Protein YceI | Periplasmic |
| Hypothetical Protein 5_*(SP5) | N/A |
| Outer Membrane Protein 2 | Transmembrane |
| Non-hemolytic phospholipase C | Periplasmic |
| DUF2147 domain-containing protein | N/A |
| Peroxiredoxin OmsC | Cytoplasmic |
| Cyclic di-GMP binding protein | Transmembrane |
| Pyrimidine-specific ribonucleoside hydrolase RihA | N/A |

| | |
|---|---|
| RidA family protein | Cytoplasmic |
| Transcription termination factor Rho | Cytoplasmic |
| Ribonuclease HII | Cytoplasmic |
| Cytochrome bd-II ubiquinol oxidase subunit 1 | N/A |
| DNA repair protein RadC | Cytoplasmic |
| Hypothetical Protein 7 | N/A |
| 6-Phosphoglucolactonase | Cytoplasmic |
| Copper resistance protein C | Periplasmic |
| Leucine--tRNA ligase | Cytoplasmic |
| 6-phosphogluconate dehydrogenase, NAD(+)-dependent, decarboxylating | Cytoplasmic |
| ATM1-type heavy metal exporter | Transmembrane |
| Rna polymerase subunit sigma | Cytoplasmic |
| Flavoheamoprotein | Cytoplasmic |
| Putative phospholipid-binding protein MlaC | Periplasmic |
| Putative membrane protein | Transmembrane |
| Putative CtpA-like serine protease | Transmembrane |
| Hypothetical Protein 8 | N/A |
| Transaldolase | Cytoplasmic |
| Conjugal transfer protein TrbF/ Type IV secretion system protein | Transmembrane |
| Membrane-bound lytic murein transglycosylase B | Transmembrane |
| Putative ABC transporter ATP-binding protein YheS | Transmembrane |

| | |
|---|---|
| KHG/KDPG aldolase | Cytoplasmic |
| Hypothetical Protein 9 | N/A |
| 6-phosphogluconate phosphatase | Cytoplasmic |
| Nucleotidyltransferase | Cytoplasmic |
| Putative deferrochelatase/peroxidase YfeX | Periplasmic |
| HAD_IIIC family phosphatase | N/A |

*Denotes the *K. rhaeticus* native SPs used in study

| Table S2. Predicted signal peptide sequences and associated translocation pathways of candidate proteins in *K. rhaeticus* | | | | | | |
|---|---|---|---|---|---|---|
| **SP** | **Associated protein_ID** | **Predicted SP sequence and associated translocase machinery** | | | | |
| | | **SP sequence** | **Sec/SP I** | **Tat/SP 1** | **Lipoprotein (Sec/SPII)** | **Other** |
| 1 | Hypothetical protein_1_Truncated endoglucanase | MGRRSFLSVMAAAG | 0.3640 | 0.6342 | 0.0009 | 0.0009 |
| 2 | Hypothetical protein_2 | MRIQIPPFRKNDPRLLLLLWSTTALFVTETCAHA | 0.3839 | 0.016 | 0.2285 | 0.3716 |
| 3 | Hypothetical protein_3 | MLMSRRLAVLALVLAPLAGLPHAAMA | 0.9388 | 0.0595 | 0.0011 | 0.0006 |
| 4 | Hypothetical protein_4 | MAKTIRSALLAALIASTPALALA | 0.9893 | 0.0044 | 0.0053 | 0.0010 |
| 5 | Hypothetical protein_5 | MPNILRRQRRQLARRPLLAGSAFMWLLSVGPAAIA | 0.2396 | 0.2274 | 0.0546 | 0.4782 |
| 6 | Putative cmcAX/ endoglucanase | MGRRSFLSVMAAAGSIPFLSTALA | 0.3640 | 0.6342 | 0.0009 | 0.0009 |
| 7 | Tolulene Transporter subunit (TodC1) | MNIMKHVLKTTSLVLCASLMAAPVVPAVFSIHAAHA | 0.9553 | 0.0119 | 0.0257 | 0.0070 |
| 8 | Virginimycin Protein (vgb) | MKTGLLSTILHRRAGAALLLSLLAPVAARA | 0.9244 | 0.0693 | 0.0027 | 0.0036 |
| 9 | Periplasmic protease | MFRDAMSDVLRSNTLTSRRFRSRLLASLVVGTVAGGCLLAAPPVRA | 0.1509 | 0.8421 | 0.0060 | 0.0009 |
| 10 | Outer membrane protein_ | MRLRAALLATSLLAAAPVAAKA | 0.9967 | 0.0016 | 0.0015 | 0.0002 |

| Table S3. Sequences of heterologous SPs used in this study, including their source organisms, associated proteins, and amino acid sequences. | | | | |
|---|---|---|---|---|
| **Name** | **Organism** | **Protein Association** | **Sequence** | **Source** |
| DsbA | *E. coli* | DsbA | MKKIWLALAGLVLAFSASA | [1] |
| PhoA | *E. coli* | PhoA | MKQSTIALALLPLLFTPVTKA | [2] |
| NSP4 | *E. coli* | Novel Signal Peptide 4 | MKKITAAAGLLLLAAQPAMA | [3] |
| YwmC | *B. subtilis* | Uncharacterized YwmC precursor | MKKRFSLIMMTGLLFGLTSPAFA | [4] |
| AmyE | *B. subtilis* | Amylase | FAKRFKTSLLPLFAGFLLLFHLVLAGPAAASA | [4] |
| EPR | *B. subtilis* | Minor Protease | MKNMSCKLVVSVTLFFSFLTIGPLAHA | [5] |
| PelB | *E. carotovora* | Periplasmic pectate lyase | MKYLLPTAAAGLLLLAAQPTMA | [6] |
| MalE | *E. coli* | Maltose binding protein | MKIKTGARILALSALTTMMFSASALA | [7] |
| OmpA | *E. coli* | Outer membrane protein A | MKKTAIAIAVALAGFATVAQA | [8] |

| Table S4. Supernatant pH, glucose consumption, and acetic acid formation after 24, 48, and 72 h of induced SP-mScarlet expression. | | | |
|---|---|---|---|
| | **Time** | | |
| **Sample** | **24h** | **48h** | **72h** |
| *Supernatant pH* | | | |
| DsbA | 6.33 | 4.85 | 4.81 |
| OmpA | 6.51 | 4.89 | 4.86 |
| SP4 | 6.53 | 4.79 | 4.80 |
| YwmC | 6.16 | 5.05 | 4.95 |
| *Glucose consumption (g $L^{-1}$)* | | | |
| DsbA | 2.8069 | 2.1463 | 2.2944 |
| OmpA | 2.4299 | 1.9768 | 2.0678 |
| SP4 | 3.0412 | 2.3268 | 2.2833 |
| YwmC | 2.4922 | 2.1642 | 2.4734 |
| *Acetic acid formation (g $L^{-1}$)* | | | |
| DsbA | 0.0239 | 0.1438 | 0.0859 |
| OmpA | 0.0256 | 0.0615 | 0.1417 |
| SP4 | 0.0314 | 0.0956 | 0.1902 |
| YwmC | 0.0426 | 0.1056 | 0.1680 |

Table S5. ProtParam predictions of molecular weights and GRAVY scores of the SPs used in this study [9].

| SP | Mw (kDa) | GRAVY score |
|---|---|---|
| DsbA | 1.99 | 1.416 |
| PhoA | 2.26 | 0.971 |
| NSP4 | 1.98 | 1.105 |
| YwmC | 2.56 | 0.857 |
| AmyE | 3.39 | 1.231 |
| EPR | 2.96 | 1.122 |
| PelB | 2.26 | 1.077 |
| MalE | 2.70 | 1.012 |
| OmpA | 2.05 | 1.295 |
| SP1 | 1.45 | 0.614 |
| SP2 | 3.95 | 0.221 |
| SP3 | 2.69 | 1.412 |
| SP4 | 2.27 | 1.361 |
| SP5 | 3.93 | 0.223 |
| SP6 | 2.45 | 0.967 |
| SP7 | 3.77 | 1.136 |
| SP8 | 3.09 | 0.920 |
| SP9 | 4.93 | 0.387 |
| SP10 | 2.18 | 1.150 |

| Table S6. KTK Entry-level parts used in the study | | |
|---|---|---|
| **KTK part** | **Description** | **Antibiotic resistance** |
| KTK_001 | Empty backbone for Entry-level parts. Contains sfGFP dropout. | Amp |
| KTK_009 | Empty backbone for D1.1. Contains LacZ dropout. | Cm |
| KTK_010 | Empty backbone for D1.2. Contains LacZ dropout. | Cm |
| KTK_016 | Empty backbone for D2.2b. Contains LacZ dropout. | Spec |
| KTK_002 | J23104 promoter | Amp |
| KTK_043 | J23101 | Amp |
| KTK_032 | pLac promoter | Amp |
| KTK_034 | pLux promoter | Amp |
| KTK_003 | RBS BBa_B0034 | Amp |
| KTK_029 | RBS BBa_B0035 | Amp |
| KTK_033 | LuxR | Amp |
| KTK_005 | Terminator sequence Bba_B0015 | Amp |
| KTK_007 | Terminator sequence Bba_B0015 | Amp |
| KTK_374 | DsbA SP | Amp |
| KTK_375 | PhoA SP | Amp |
| KTK_376 | NSP4 SP | Amp |
| KTK_377 | YwmC SP | Amp |
| KTK_378 | EPR SP | Amp |
| KTK_379 | SP1 SP | Amp |
| KTK_380 | SP2 SP | Amp |
| KTK_381 | SP3 SP | Amp |
| KTK_382 | SP4 SP | Amp |
| KTK_383 | SP5 SP | Amp |
| KTK_384 | SP6 SP | Amp |
| KTK_385 | SP7 SP | Amp |
| KTK_386 | SP9 SP | Amp |
| KTK_387 | SP10 SP | Amp |
| KTK_389 | PelB SP | Amp |
| KTK_251 | mScarlet used to make 1.3 mScarlet with a His tag | Amp |
| KTK_252 | mScarlet used to make 1.3b mScarlet with a His tag | Amp |
| KTK_390 | β-lactamase 1.3b, to construct SP-fusion reporter | Amp |
| KTK_391 | β-lactamase 1.3 | Amp |

Table S7. List of expression plasmids used in this study. Constitutively expressed fusion proteins were constructed with Komagataeibacter Toolkit as level 1 (D1.1) plasmids in the KTK_010 vector (CmR). Inducibly expressed fusion proteins were constructed with KTK as level 2 (D2.2) plasmids in the KTK_016 vector (Spec).

| **Expressed construct** | **Plasmid parts** |
|---|---|
| D1.1 Bla | J23101_003-RBS_GFP_Bla_BBa_B0015 |
| D1.1 GFP | J23101_003-RBS_GFP_sfGFP_BBa_B0015 |
| D1.1 DsbA_Bla | J23101_003-RBS_GFP_DsbSS_Blac_BBa_B0015 |
| D1.1 PhoAss_Bla | J23101_003-RBS_GFP_PhoAss_Blac_BBa_B0015 |
| D1.1 NSP4_Bla | J23101_003-RBS_GFP_NSP4_Blac_BBa_B0015 |
| D1.1 YwmC_Bla | J23101_003-RBS_GFP_YwmC_Blac_BBa_B0015 |
| D1.1 AmyESP_Bla | J23101_003-RBS_GFP_AmyESP_Blac_BBa_B0015 |
| D1.1 EPR_Bla | J23101_003-RBS_GFP_EPR_Bla_BBa_B0015 |
| D1.1 SP1_Bla | J23101_003-RBS_GFP_SP1_Bla_BBa_B0015 |
| D1.1 SP2_Bla | J23101_003-RBS_GFP_SP2_Bla_BBa_B0015 |
| D1.1 SP3_Bla | J23101_003-RBS_GFP_SP3_Bla_BBa_B0015 |
| D1.1 SP4_Bla | J23101_003-RBS_GFP_SP4_Bla_BBa_B0015 |
| D1.1 SP5_Bla | J23101_003-RBS_GFP_SP5_Bla_BBa_B0015 |
| D1.1 SP6_Bla | J23101_003-RBS_GFP_SP6_Bla_BBa_B0015 |
| D1.1 SP7_Bla | J23101_003-RBS_GFP_SP7_Bla_BBa_B0015 |
| D1.1 SP8_Bla | J23101_003-RBS_GFP_SP8_Bla_BBa_B0015 |
| D1.1 SP9_Bla | J23101_003-RBS_GFP _SP9_Bla_BBa_B0015 |
| D1.1 SP10_Bla | J23101_003-RBS_GFP _SP10_Bla_BBa_B0015 |
| D1.1 Bla | J23104_003-RBS_GFP_Bla_BBa_B0015 |
| D1.1 YwmC_Bla | J23104_003-RBS_GFP_YwmC_Bla_BBa_B0015 |
| D1.1 EPR_Bla | J23104_003-RBS_GFP_EPR_Bla_BBa_B0015 |
| D1.1 mScarlet | J23104_029-RBS_ mScarlet-his_BBa_B0015 |
| D1.1 DsbA-mScarlet-his | J23104_029-RBS_DsbA_mScarlet_BBa_B0015 |
| D1.1 MalE-mScarlet-his | J23104_029-RBS_MalE_mScarlet_BBa_B0015 |
| D1.1 OmpA-mScarlet-his | J23104_029-RBS_OmpA_mScarlet_BBa_B0015 |
| D1.1 PelB-mScarlet-his | J23104_029-RBS_PelB_mScarlet-his_BBa_B0015 |
| D1.1 SP2-mScarlet-his | J23104_029-RBS_SP2_mScarlet-his_BBa_B0015 |
| D1.1 SP4-mScarlet-his | J23104_029-RBS_SP4_mScarlet-his_BBa_B0015 |
| D1.1 SP10-mScarlet-his | J23104_029-RBS_SP10_mScarlet-his_BBa_B0015 |
| D1.1 YwmC-mScarlet-his | pLac_029-RBS_YwmC_mScarlet-his_BBa_B0015 |
| D2.2 mScarlet | J23104_003-RBS_LuxR_BBa_B0015_pLux_029-RBS_mScarlet-his_BBa_B0015 |
| D2.2 LuxR_DsbA-mScarlet | J23104_003-RBS_LuxR_BBa_B0015_pLux_029-RBS_DsbA_mScarlet-his_BBa_B0015 |
| D2.2 LuxR_MalE-mScarlet | J23104_003-RBS_LuxR_BBa_B0015_pLux_029-RBS_MalE_mScarlet-his_BBa_B0015 |
| D2.2 LuxR_OmpA-mScarlet | J23104_003-RBS_LuxR_BBa_B0015_pLux_029-RBS_OmpA_mScarlet-his_BBa_B0015 |
| D2.2 LuxR_PelB-mScarlet | J23104_003-RBS_LuxR_BBa_B0015_pLux_029-RBS_PelB_mScarlet-his_BBa_B0015 |

| | |
|---|---|
| D2.2 LuxR_SP2-mScarlet | J23104_003-RBS_LuxR_BBa_B0015_pLux_029-RBS_SP2_mScarlet-his_BBa_B0015 |
| D2.2 LuxR_SP4-mScarlet | J23104_003-RBS_LuxR_BBa_B0015_pLux_029-RBS_SP4_mScarlet-his_BBa_B0015 |
| D2.2 LuxR_SP10-mScarlet | J23104_003-RBS_LuxR_BBa_B0015_pLux_029-RBS_SP10_mScarlet-his_BBa_B0015 |
| D2.2 LuxR_YwmC-mScarlet | J23104_003-RBS_LuxR_BBa_B0015_pLux_029-RBS_YwmC_mScarlet-his_BBa_B0015 |

| Table S8. List of primers used in the study | | | |
|---|---|---|---|
| **Primer name** | **Description** | **Sequence** | **Reference** |
| KTK_PelB_Frd_1.3a | To create 1.3a entry-level of PelB | CGTCTCCTCGGTCTCCTATGAAATATCTGCTGCCGACCGCA | This study |
| KTK_PelB_Rev_1.3a | To create 1.3a entry-level of PelB | CGTCTCCGGTCTCAAGAACCTGCCATGGTCGGCTGTGC | This study |
| mScarletHis_KTK_EL_1.3b_FOR | To create 1.3b entry-level of mScarlet | CGTCTCCTCGGTCTCATTCTATGGTCAGTAAAGGCGAAGCAGTTATCAAAG | This study |
| mScarletHis_KTK_EL_1.3b_REV | To create 1.3b entry-level of mScarlet with C-terminal His-tag | CGTCTCGGGTCTCTTGACTTATTAATGGTGGTGATGGTGATGGTGAGCAGCGGCCTTGTATAACTCGTCCATACCGCCCG | This study |
| JV_mScarlet1.3_EL_FOR | To create 1.3 entry-level of mscarlet | CGT CTC CTC GGT CTC CTA TGG TCA GTA AAG GCG AAG CAG | This study |
| KTK_seq_FP | Sequencing primer for Entry-level cloning flanking left of the insertion | TATAGTCCTGTCGGGTTTCGCC | [10] |
| KTK_Seq_RP | Sequencing primer for Entry-level cloning flanking right of the insertion | CCGGTGAGCGTGGGTCCCGCGGTATC | [10] |
| KTKStdFrdD1_D2 | Primer for region left of insertion in KTK_D1 and KTK_D2 constructs. | AGGGCGGCGGATTTGTCC | [10] |
| PQE30Rev | Primer for region right of insertion in KTK_D1 and KTK_D2 constructs | GTTCTGAGGTCATTACTGG | [10] |
